# LeDNA: a cut-and-build toolkit to democratize education on CRISPR gene editing technology

**DOI:** 10.1101/2024.10.09.617025

**Authors:** Guilherme E. Kundlatsch, Alina S. L. Rodrigues, Vitória F. B. Zocca, Laura A. S. Amorim, Gabriela B. de Paiva, Almiro P. S. Neto, Juliana A. D. B. Campos, Danielle B. Pedrolli

## Abstract

We introduce LeDNA, a versatile educational toolkit designed for teaching fundamental genetics and CRISPR-Cas gene editing principles in diverse settings, regardless of existing infrastructure. Fabricated using laser-cutting techniques, LeDNA is an open-source resource suitable for students across educational levels, from high school to graduate studies. Given the transformative potential of CRISPR technology in various fields, including medicine and agriculture, a widespread understanding of its principles is essential for informed public discourse and acceptance. By providing a readily accessible and affordable tool, LeDNA aims to democratize genetics and CRISPR education globally, fostering a more informed and engaged community.

## INTRODUCTION

The limited effectiveness of a lecture-based biology class system had already been demonstrated in the first half of the last century^1–3^. Several authors have reported better student learning outcomes accomplished by active learning methodologies in recent years ^4– 6^. Therefore, it is not surprising that active learning methodologies using diverse model organisms have been reported since the paradigm-shifting implementation of the CRISPR-Cas gene editing technology (Table S1). Most of these works, however, focused on undergraduate students. Considering the widespread impact that CRISPR-Cas will have on society in the next decades, we developed a new open-access active-learning methodology to teach basic genetics and gene editing in secondary schools, aiming at a broader audience.

Not all students have access to a science laboratory in secondary schools. For example, less than half of secondary schools in Brazil have such facilities^7^. Furthermore, the different legislations around the globe for manipulating GMOs may hamper or even render the use of experimental-based approaches impossible in many institutions. An alternative would be the use of computer simulations, which have been reported to positively impact teaching biotechnology topics^8–11^. Recently, Pal et al.^12^ reported the development of a virtual teaching module for CRISPR-Cas. However, these approaches also come up against infrastructure limitations. As an example, 69.8% of schools in Brazil lack dedicated computer laboratories (Figure S1). In such settings, the only remaining alternative may be noncomputer educational games, which do not require specific infrastructure and have been successfully applied to teach several biology topics (Table S2). Such an approach for CRISPR-Cas teaching has not yet been reported.

Inspired by the proliferation of maker spaces and digital fabrication equipment in educational institutions^14^, we designed LeDNA, an open-access tool fabricated through laser-cutting. LeDNA allows users to design DNA sequences, translate and transcribe them, as well as edit them reproducing the CRISPR-Cas mechanism. The production of LeDNA costs less than 10 US Dollars per kit and the files are available in five languages (Supplementary Material). Its operation demands no laboratory infrastructure, and the user-friendly open-access files empower any institution with a laser cutter to swiftly produce it locally. Additionally, we developed a streamlined experimental laboratory class using *Bacillus subtilis* (an FDA’s GRAS bacterium) and a theoretical class. Finally, we applied the three interventions sequentially in a Brazilian secondary school with low-income students. We also developed a new tool to assess the learning of not only the technical aspects of CRISPR-Cas but also the understanding of both basic genetics and the applications of gene editing in daily life, expanding the assessment tools described by previous authors^15–20^.

## RESULTS

### LeDNA design

To design LeDNA, we established twelve requirements to be met to prevent users from violating the central dogma of molecular biology and to allow freedom of choice for constructing the DNA sequence and editing. A solution was developed for each criterion (Table 1). LeDNA is composed of 186 pieces representing the sugar-phosphate backbone of DNA and RNA, DNA and RNA nucleotides, ribosomes, amino acids, hydrogen bonds, peptide bonds, and the Cas protein (Figures S2-S10). The user can assemble a double-strand DNA and perform transcription and translation, as well as CRISPR-Cas gene editing. Assembling any 12-nucleotide DNA sequence of choice is only possible when using canonical Watson–Crick base pairing. The pieces were designed to permit only the assembly of the corresponding RNA and protein based on the user-defined DNA sequence. Finally, the user can edit the original sequence through the pieces representing CRISPR-Cas to create a new RNA transcript and the corresponding protein sequence. Each kit was designed to be collaboratively used by 4 students in 90-minute classes (Figure 1).

**Table 1.**
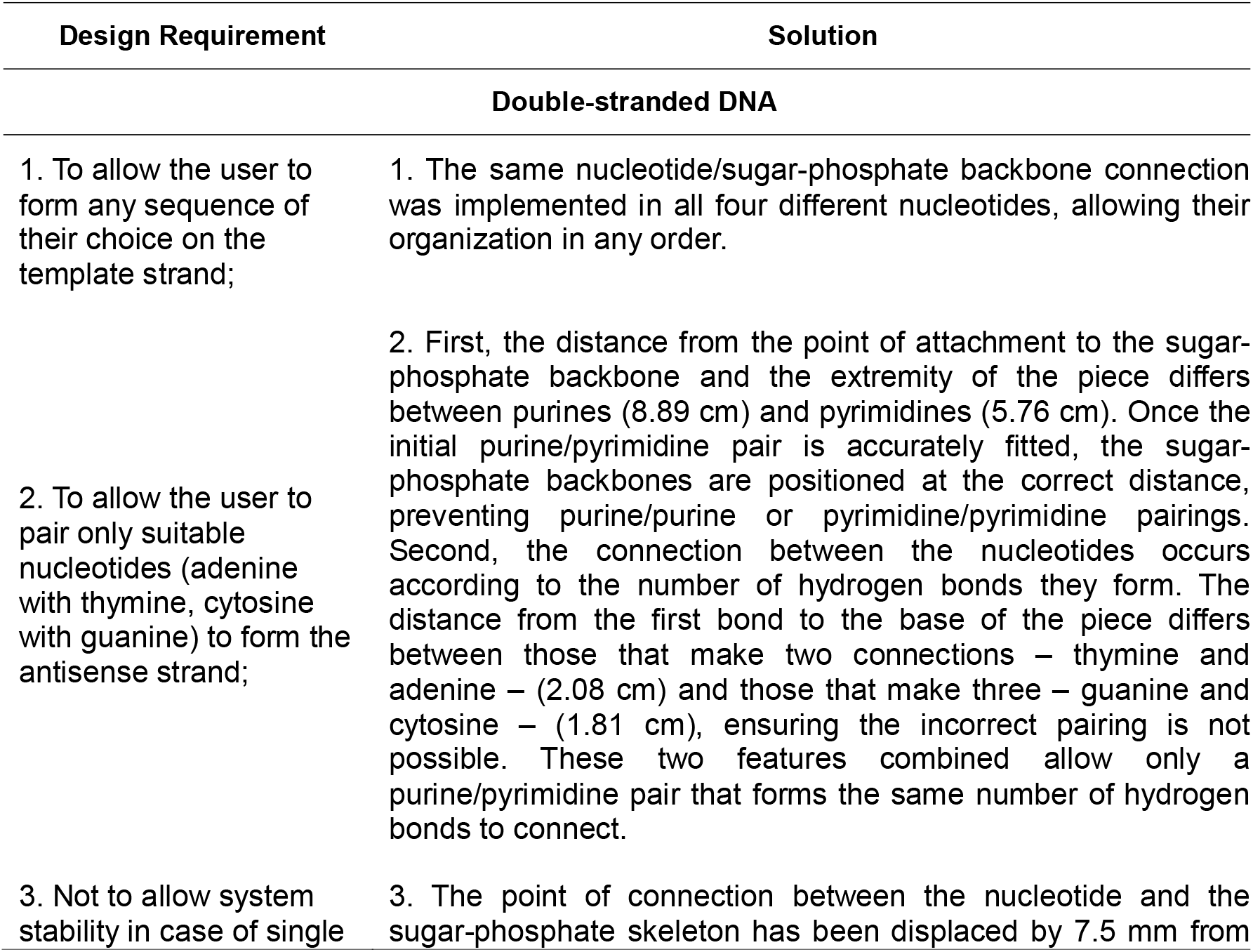
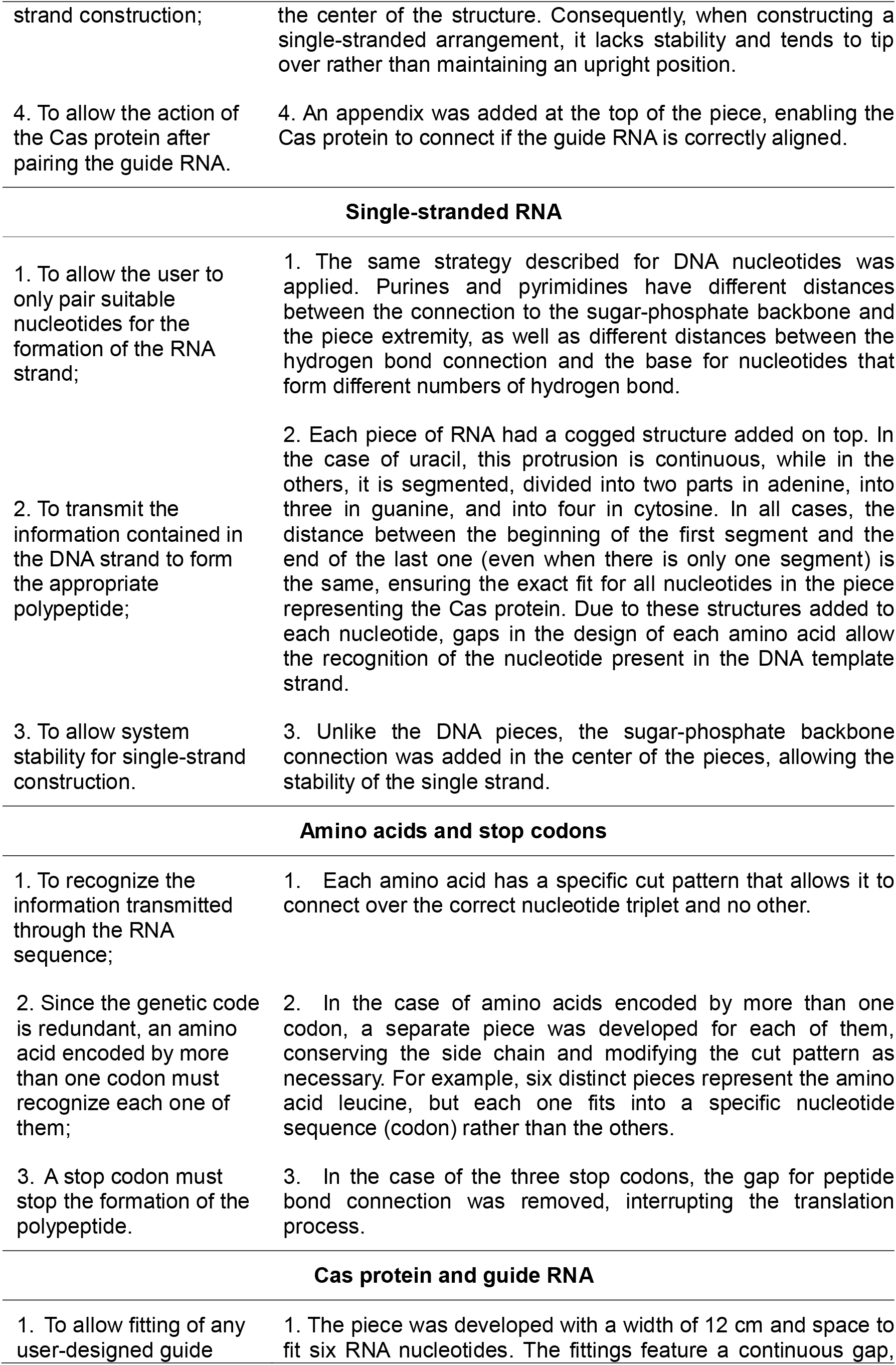
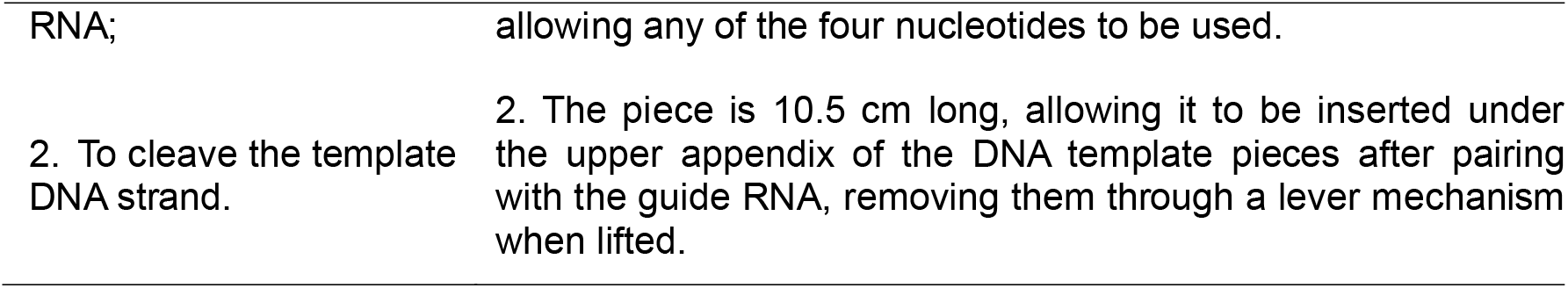
Design requirements for LeDNA Design.

**Figure 1.**
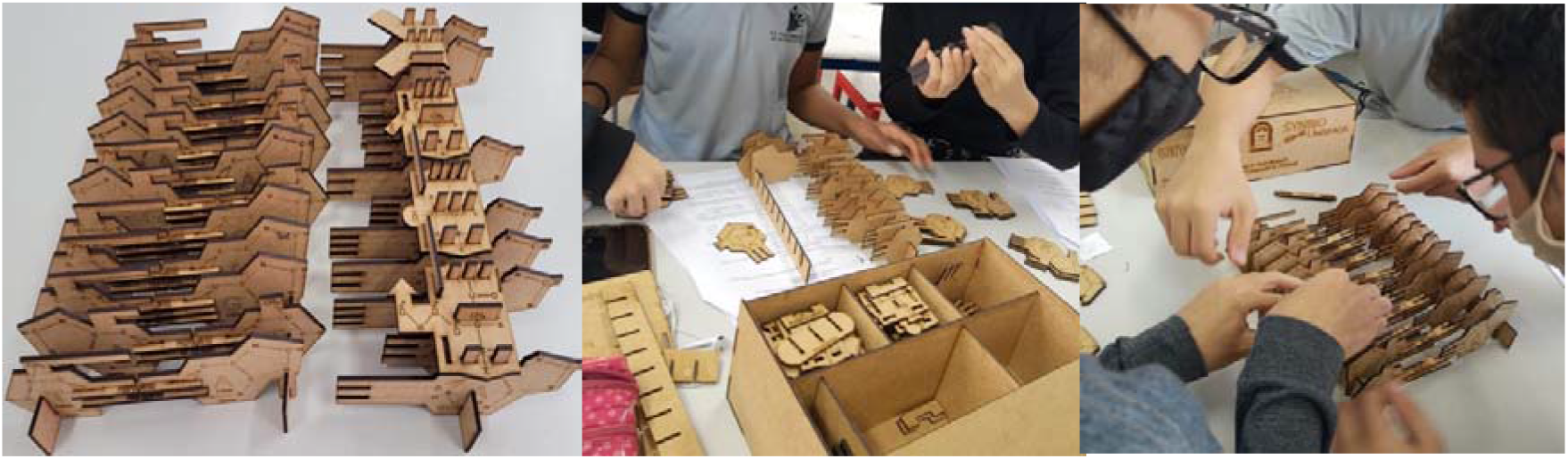
The LeDNA toolkit is designed to be used collaboratively by students. It is composed of 186 pieces representing the sugar-phosphate backbone of DNA and RNA, DNA and RNA nucleotides, ribosomes, amino acids, hydrogen bonds, peptide bonds, and the Cas protein.

### Laboratory class design

To compare different learning activities, we established a simplified editing protocol tailored to the prevalent limitations of school infrastructure and constrained class time. The CRISPR-Cas gene editing methodology for *B. subtilis* described by Altenbuchner^21^ was successfully adapted into a 90-minute session, reducing the original duration of several days. This was accomplished by reducing the cell regeneration time after transformation to 45 min and simulating the plasmid loss process in the meantime (Figure 2). The equipment requirements were kept minimal, the water bath for thawing frozen competent cells at 37°C was replaced by participants holding samples in their hands tightly, Drigalski spatulas were substituted by swirling Petri dishes over a table, and all incubations were carried out at room temperature. Despite the absence of sterilization methods, the majority of transformed plates were not contaminated, and the shortened transformation method resulted in some transformants on plates (Figure S11).

**Figure 2.**
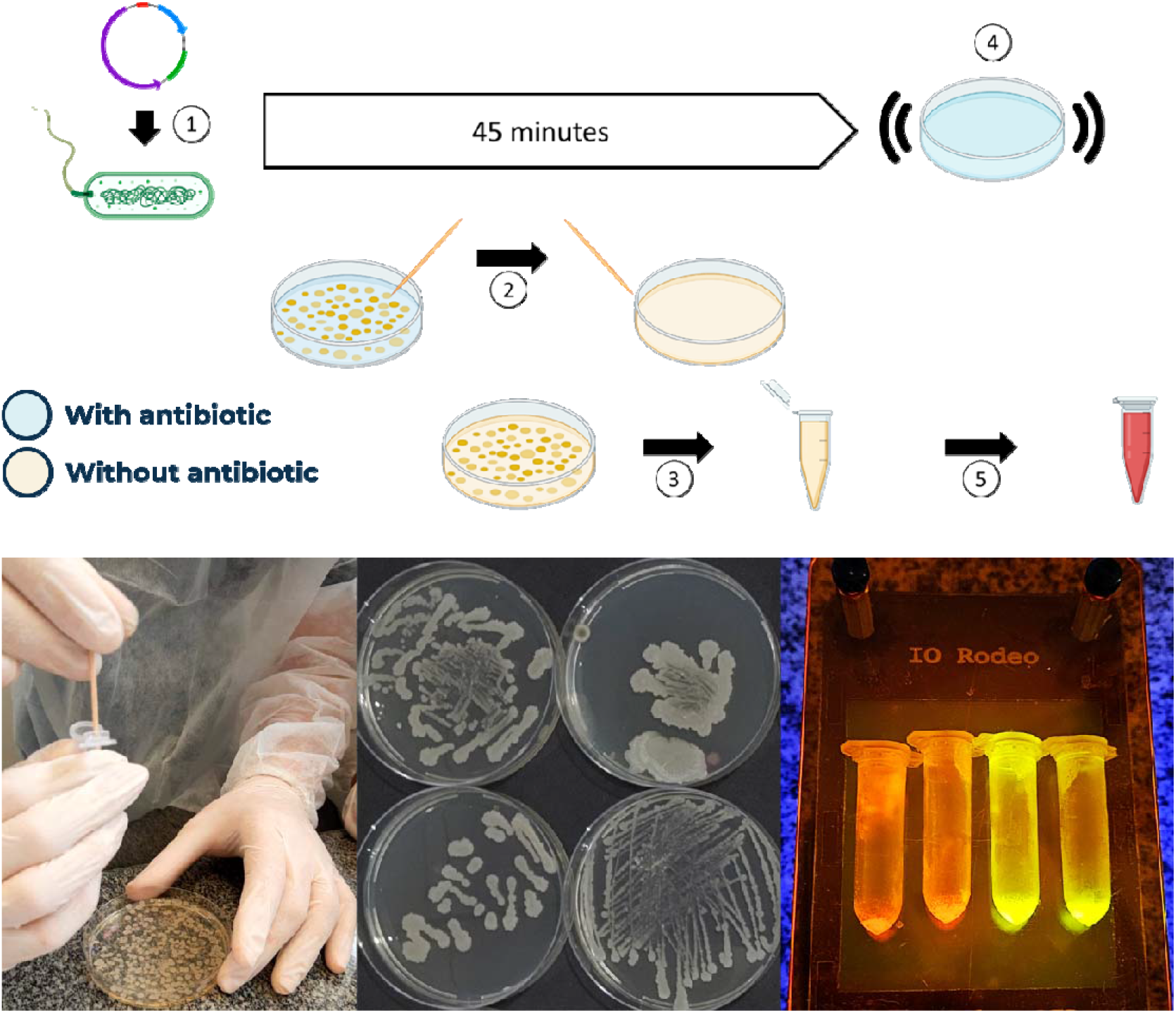
Simplified CRISPR-Cas protocol for wet lab lecture. After transformation (1), cells were incubated on the bench. Meanwhile, the participants received pre-transformed cells and streaked them on an antibiotic-free medium to simulate the plasmid loss process (2). Subsequently, they received a new Petri dish containing cells without plasmid and inoculated them in a liquid medium (3). Next, the students plated the cells they had transformed in the first step (4). Finally, the students visualized the fluorescence of cells producing either RFP or GFP using a portable transilluminator.

### Assessments

Students’ demographic profile was assessed once before the interventions (Tables S3 and S4). Students’ attitudes towards science, assessed before and after each intervention, did not change significantly throughout the study (Table S5). The students’ specific knowledge was also assessed before and after each intervention using an evaluation questionnaire. The questionnaire comprises nine questions to evaluate knowledge in Basic Genetics, CRISPR/Cas Mechanism, and Gene Editing Applications. The questionnaire was validated with the answers from students who participated in all activities, totaling 224 responses. The frequency of responses (Table S6), difficulty, and discrimination of each item (Table S7) were as expected. As expected, the questions in the first session of Basic Genetics resulted in a higher frequency of correct answers than the other two, since they assessed topics of the high school curriculum. All nine questions fit a two-parameter logistic (2PL) item response theory model (Table S8). The alpha coefficient (Cronbach’s Alpha) was calculated as 0.8464, indicating the test’s reliability.

### Impact of the 3-Stages Teaching Approach on High School Students

To compare different teaching methods and their combination, we implemented a 3-Stages Teaching Approach encompassing the LeDNA, wet lab, and theoretical lectures. Each intervention lasted 90 minutes and was applied at one-week intervals. The three interventions were evaluated both individually and combined in different orders (Figure 3).

**Figure 3.**
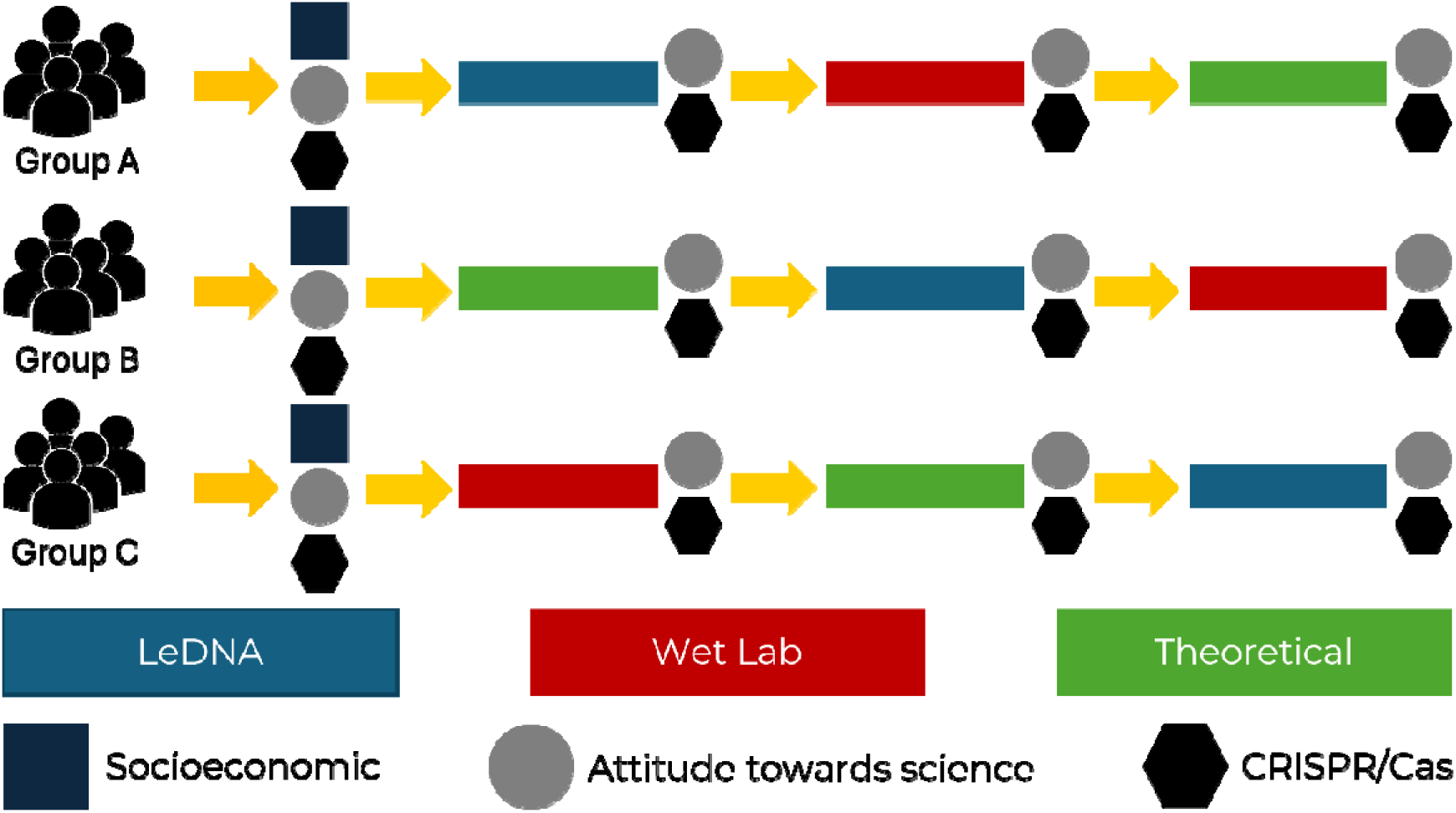
The 3-Stages Teaching Approach. Each group of students was submitted to all three teaching methodologies but in different orders. Students’ demographic profile was assessed once before the interventions. Attitudes toward science and specific knowledge were assessed before the interventions and after each stage.

All groups commenced from approximate initial scores. Each isolated intervention positively influenced participants’ knowledge of CRISPR-Cas, with all groups achieving significantly higher scores after the initial intervention compared to their baseline scores. Subsequent activities led to a further increase in students’ performance (Table 2). It is clear that the repetition of the interventions had a positive impact on the participants’ learning. Nevertheless, the uneven score gain after each stage for each group suggests the nature of the activity influenced the knowledge acquisition. Groups A and C had higher score improvement following the LeDNA activity delivered as the first and third lessons, respectively. The wet lab activity resulted in the second-highest score gain for all three groups regardless of the order in which it was delivered. The results demonstrate the effectiveness of all three methodologies and showcase their complementary effect. Moreover, they indicate that LeDNA is an efficient tool suitable for standalone use or in conjunction with one or two other activities, depending on the available infrastructure.

**Table 2.** High School students’ performance before and after each intervention.

| Complete Learning Assessment |  |  |  |  |  |  |
| --- | --- | --- | --- | --- | --- | --- |
| Group | Score | Before | Stage 1 | Stage 2 | Stage 3 | Total gain |
| A | Mean ± SD | 0.997±1.305<sup>a, A</sup> | 4.180±1.726<sup>b, B</sup> | 6.643±1.752<sup>b, C</sup> | 7.512±1.430<sup>b, C</sup> | 6.515 |
|  | Gain (%) | 0 | 48.9 | 37.8 | 13.3 | 100 |
| B | Mean ± SD | 0.955±1.452<sup>a, A</sup> | 3.907±1.671<sup>b, B</sup> | 4.283±1.429<sup>a, B</sup> | 6.644±1.818<sup>ab, C</sup> | 5.689 |
|  | Gain (%) | 0 | 51.9 | 6.6 | 41.5 | 100 |
| C | Mean ± SD | 0.994±0.938<sup>a, A</sup> | 2.131±1.129<sup>a, AB</sup> | 2.955±2.588<sup>a, B</sup> | 5.120±1.795<sup>a, C</sup> | 4.126 |
|  | Gain (%) | 0 | 27.6 | 20.0 | 52.5 | 100 |

■ LeDNA ■ Wet Lab ■ Theoretical
Uppercase letters indicate significant statistical differences between columns, while lowercase letters indicate significant statistical differences between rows.

### Learning per session

Students were assessed on three CRISPR technology-related topics: Basic Genetics, CRISPR-Cas Mechanism, and Gene Editing Applications. Unexpectedly, high-school students scored poorly in Basic Genetics in the initial assessment (Table 3), despite this session evaluating topics students should have previously learned according to the national curriculum. We speculate that the poor performance was due to educational difficulties arising from the pandemic, especially in public education institutions in Brazil^22^. At the end of the three activities, all groups reached a high score in this session, indicating that the interventions were efficient in reversing the adverse scenario.

**Table 3.**
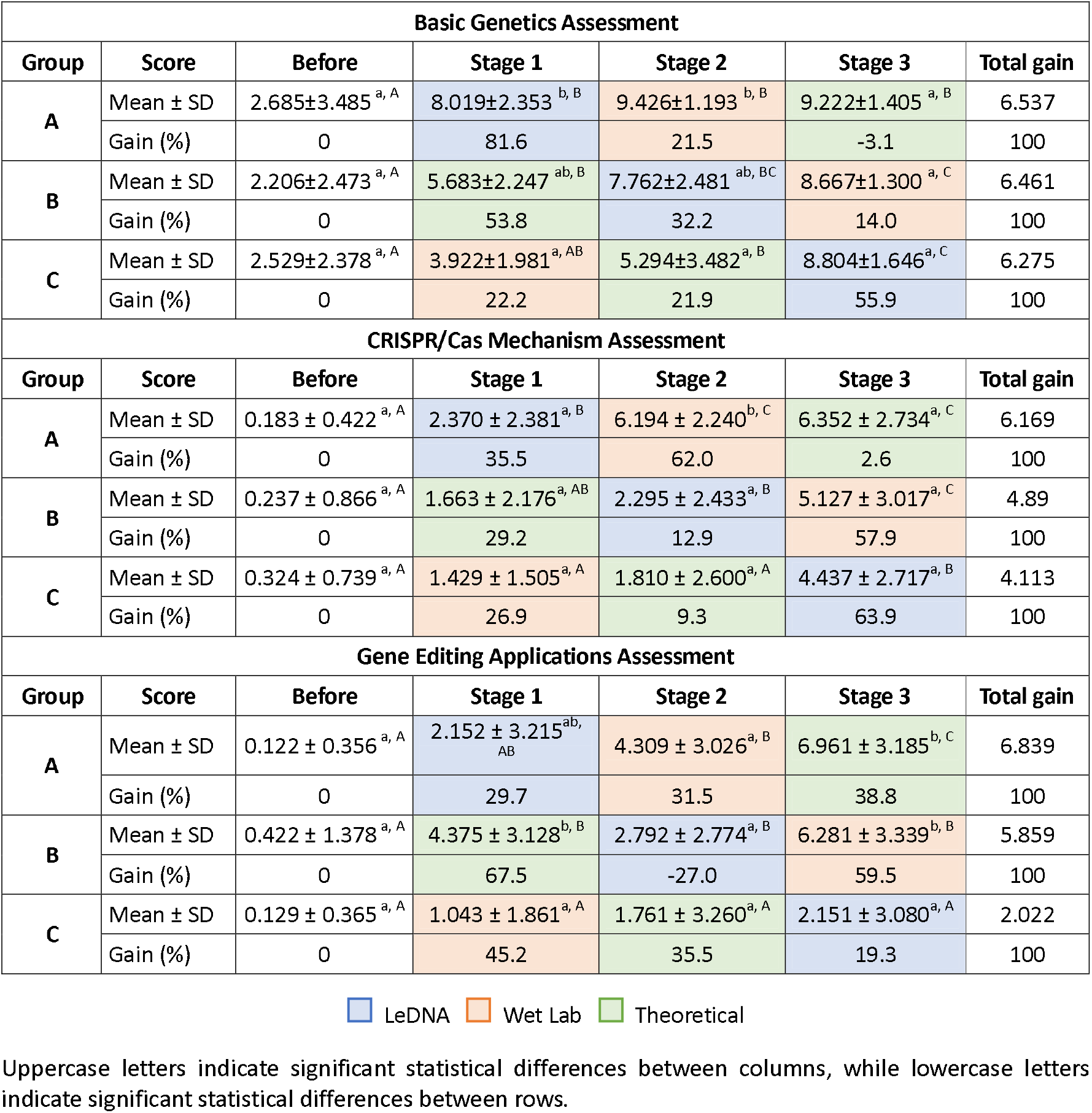
High School students’ performance per session.

| Basic Genetics Assessment |  |  |  |  |  |  |
| --- | --- | --- | --- | --- | --- | --- |
| Group | Score | Before | Stage 1 | Stage 2 | Stage 3 | Total gain |
| A | Mean ± SD | 2.685±3.485<sup>a, A</sup> | 8.019±2.353<sup>b, B</sup> | 9.426±1.193<sup>b, B</sup> | 9.222±1.405<sup>a, B</sup> | 6.537 |
|  | Gain (%) | 0 | 81.6 | 21.5 | -3.1 | 100 |
| B | Mean ± SD | 2.206±2.473<sup>a, A</sup> | 5.683±2.247<sup>ab, B</sup> | 7.762±2.481<sup>ab, BC</sup> | 8.667±1.300<sup>a, C</sup> | 6.461 |
|  | Gain (%) | 0 | 53.8 | 32.2 | 14.0 | 100 |
| C | Mean ± SD | 2.529±2.378<sup>a, A</sup> | 3.922±1.981<sup>a, AB</sup> | 5.294±3.482<sup>a, B</sup> | 8.804±1.646<sup>a, C</sup> | 6.275 |
|  | Gain (%) | 0 | 22.2 | 21.9 | 55.9 | 100 |
| CRISPR/Cas Mechanism Assessment |  |  |  |  |  |  |
| Group | Score | Before | Stage 1 | Stage 2 | Stage 3 | Total gain |
| A | Mean ± SD | 0.183 ± 0.422<sup>a, A</sup> | 2.370 ± 2.381<sup>a, B</sup> | 6.194 ± 2.240<sup>b, C</sup> | 6.352 ± 2.734<sup>a, C</sup> | 6.169 |
|  | Gain (%) | 0 | 35.5 | 62.0 | 2.6 | 100 |
| B | Mean ± SD | 0.237 ± 0.866<sup>a, A</sup> | 1.663 ± 2.176<sup>a, AB</sup> | 2.295 ± 2.433<sup>a, B</sup> | 5.127 ± 3.017<sup>a, C</sup> | 4.89 |
|  | Gain (%) | 0 | 29.2 | 12.9 | 57.9 | 100 |
| C | Mean ± SD | 0.324 ± 0.739<sup>a, A</sup> | 1.429 ± 1.505<sup>a, A</sup> | 1.810 ± 2.600<sup>a, A</sup> | 4.437 ± 2.717<sup>a, B</sup> | 4.113 |
|  | Gain (%) | 0 | 26.9 | 9.3 | 63.9 | 100 |
| Gene Editing Applications Assessment |  |  |  |  |  |  |
| Group | Score | Before | Stage 1 | Stage 2 | Stage 3 | Total gain |
| A | Mean ± SD | 0.122 ± 0.356<sup>a, A</sup> | 2.152 ± 3.215<sup>ab, AB</sup> | 4.309 ± 3.026<sup>a, B</sup> | 6.961 ± 3.185<sup>b, C</sup> | 6.839 |
|  | Gain (%) | 0 | 29.7 | 31.5 | 38.8 | 100 |
| B | Mean ± SD | 0.422 ± 1.378<sup>a, A</sup> | 4.375 ± 3.128<sup>b, B</sup> | 2.792 ± 2.774<sup>a, B</sup> | 6.281 ± 3.339<sup>b, B</sup> | 5.859 |
|  | Gain (%) | 0 | 67.5 | -27.0 | 59.5 | 100 |
| C | Mean ± SD | 0.129 ± 0.365<sup>a, A</sup> | 1.043 ± 1.861<sup>a, A</sup> | 1.761 ± 3.260<sup>a, A</sup> | 2.151 ± 3.080<sup>a, A</sup> | 2.022 |
|  | Gain (%) | 0 | 45.2 | 35.5 | 19.3 | 100 |

■ LeDNA ■ Wet Lab ■ Theoretical
Uppercase letters indicate significant statistical differences between columns, while lowercase letters indicate significant statistical differences between rows.

Although LeDNA was developed to teach CRISPR-Cas gene editing, the results demonstrate that it is a useful tool for teaching basic genetics both independently and complementarily to other approaches. When evaluating the learning per session, LeDNA yielded the higher score gain from all interventions as first activity for Group A (81%) in basic genetics (Table 3). Moreover, as the third activity for Group C, LeDNA was responsible for the highest score improvement (56%) in basic genetics for this group. Even for Group B, which had a great gain in score after the theoretical lecture, LeDNA as a second activity improved the students’ score significantly (32%). Together, the theoretical lecture followed by the LeDNA session were responsible for 86% of the final score gain in Group B, demonstrating the potential of LeDNA as a complementary tool in the classroom. By allowing students to interact with and visualize the structure of biomolecules such as DNA, RNA, and proteins, and to replicate the central dogma of biology, LeDNA brings the microscopic world of molecular biology to our macroscopic universe, thereby facilitating the learning process.

Since CRISPR-Cas is not a topic currently addressed in the high school curriculum, it was not a surprise that most students scored zero in the initial assessment in the CRISPR-Cas Mechanism session (Table 3). There was no significant difference between the groups after the implementation of the three activities, and the final score in every group was significantly higher than the initial score. However, it is interesting to note that all groups reached a statistically distinct score from the initial score only after the LeDNA session: Group A after the first activity, Group B after the second, and Group C after the third. Once again, the results highlight what a powerful tool LeDNA is, enabling students to better understand the intricacies of gene editing.

Unlike the previous learning sessions, in the Gene Editing Applications segment the theoretical lecture resulted in the most significant score improvements. Starting with the theoretical lecture, Group B achieved a 67% score gain after this single activity. A similar score gain (70%) was reached by Group A only after a combination of the wet lab followed by the theoretical lecture as second and third activities. Starting with the wet lab activity, Group C did not reach any significant score improvement throughout the 3-Stage process. One possible explanation for this phenomenon is that without comprehending what they were performing during the wet lab procedure, the students did not have their interest in the technique’s real-life applications sparked to the same extent as the other two groups. This outcome highlights that the combination of various activities enhances participants’ learning. Therefore, we suggest that in educational settings with infrastructure for conducting a practical lesson, LeDNA can be integrated as a preamble to enhance the effectiveness of the wet lab activity.

To provide anecdotal insight into the participants’ perception of CRISPR-Cas after the activities, some of the participants’ open responses regarding the future applications of the technology are listed in Table S9.

### Impact of the 3-Stages Teaching Approach on Undergraduate and Graduate Students

Although LeDNA and the 3-Stages Teaching Approach have been conceived for use in high school education, we found out that they are useful for teaching CRISPR-Cas Gene Editing to undergrad and graduate students. Since they already started with a more advanced understanding of Basic Genetics and Gene Editing Applications, their total score gain was lower than that of high school students. The first activity was already effective in improving their score, regardless of the methodology used (Table 4). As a single intervention, the theoretical lecture resulted in the highest score gain. From the three learning sessions evaluated, we only detected significant score improvement in the CRISPR-Cas Mechanism segment for undergrad and graduate students (Table 4). Again, any first intervention resulted in score improvement, regardless of which one. Interestingly, the LeDNA and the wet lab activities were equivalent in knowledge gain as the first activity, demonstrating that LeDNA may be efficiently used as a replacement for wet lab classes. Moreover, LeDNA as the last intervention for Group C was effective in improving the students’ scores in the CRISPR-Cas Mechanism session to a higher level than the other two groups. This result highlights the effectiveness of LeDNA, whether used alone or alongside other activities, in conveying the intricacies of gene editing machinery.

**Table 4.** Undergraduate and Graduate students’ performance after each intervention.

| Total score |  |  |  |  |  |  |
| --- | --- | --- | --- | --- | --- | --- |
| Group | Score | T0 | T1 | T2 | T3 | Total gain |
| A | Mean ± SD | 3.795 ± 1.210<sup>a, A</sup> | 5.469 ± 1.437<sup>a, B</sup> | 6.382 ± 1.497<sup>a, BC</sup> | 6.501 ± 1.382<sup>a, C</sup> | 2.706 |
|  | Gain (%) | 0 | 61.9 | 33.7 | 4.4 | 100 |
| B | Mean ± SD | 4.061 ± 1.536<sup>a, A</sup> | 6.252 ± 1.929<sup>a, B</sup> | 6.149 ± 1.755<sup>a, B</sup> | 6.653 ± 1.792<sup>a, B</sup> | 2.592 |
|  | Gain (%) | 0 | 84.5 | -4.0 | 19.4 | 100 |
| C | Mean ± SD | 3.790 ± 2.217<sup>a, A</sup> | 5.624 ± 1.776<sup>a, B</sup> | 6.791 ± 1.254<sup>a, BC</sup> | 7.830 ± 1.256<sup>a, C</sup> | 4.040 |
|  | Gain (%) | 0 | 45.4 | 28.9 | 25.7 | 100 |
| CRISPR/Cas Mechanism |  |  |  |  |  |  |
| Group | Score | T0 | T1 | T2 | T3 | Total gain |
| A | Mean ± SD | 0.420±1.048<sup>a, A</sup> | 3.471±2.647<sup>a, B</sup> | 4.369±3.052<sup>a, B</sup> | 4.539±2.945<sup>a, B</sup> | 6.169 |
|  | Gain (%) | 0 | 74.1 | 21.8 | 4.1 | 100 |
| B | Mean ± SD | 1.660±2.080<sup>a, A</sup> | 5.837±2.029<sup>b, B</sup> | 4.463±2.584<sup>a, B</sup> | 5.127±2.058<sup>a, B</sup> | 4.89 |
|  | Gain (%) | 0 | 120.5 | -39.6 | 19.2 | 100 |
| C | Mean ± SD | 0.398±0.776<sup>a, A</sup> | 3.469±2.621<sup>a, B</sup> | 4.509±2.145<sup>a, B</sup> | 7.953±2.200<sup>b, C</sup> | 4.113 |
|  | Gain (%) | 0 | 40.6 | 13.8 | 45.6 | 100 |

■ LeDNA ■ Wet Lab ■ Theoretical
Uppercase letters indicate significant statistical differences between columns, while lowercase letters indicate significant statistical differences between rows.

## DISCUSSION

The CRISPR-Cas gene editing technology has been a catalyst for unprecedented advancements in medicine, agriculture, and biotechnology. LeDNA brings this revolutionary technology to a broader public, enabling students to physically manipulate and explore the usually invisible molecular machinery.

Unlike the vast majority of other tools developed for CRISPR-Cas education (Table S1), LeDNA does not require any laboratory infrastructure. Although we were able to adapt a wet lab protocol to be used in high school lectures with limited laboratory facilities, the process still requires pre-work in a structured lab for competent cell preparation and proper inactivation and disposal of contaminated material. A cell-free alternative, Biobits™ Health^23^, partially bypasses infrastructure limitations but still requires an incubator and an imager while not allowing direct visualization of the CRISPR-Cas molecular mechanisms. In another approach, to overcome infrastructure limitations during the pandemic, “How to grow (almost) anything”^24^ created a distributed network model for distance-based laboratory learning for synthetic biology. The course applied student-run home experiments, cloud simulations, and both robot-assisted and teaching-assisted remote experiments. Despite bypassing local infrastructure restrictions, this resource-intensive approach may not be feasible to be ubiquitously implemented in many countries.

In an approach similar to LeDNA, SynBio in 3D^25^ proposed a decentralized production of 3D-printed pieces to visualize molecular mechanisms. The distinction from our work lies in its more narrowly defined subject matter, using a genetic toggle switch as a proof of concept, and its application to a cohort of students with a higher level of specialization.

Although comparing the learning gains from using LeDNA with other methodologies is not possible due to different assessment instruments, we demonstrate that the use of this new tool both individually and in conjunction with theoretical and wet lab lectures results in significant gains in knowledge acquisition. LeDNA stands out from previously described methodologies due to its low production cost, achieved by using MDF as raw material. Each kit costs 10 US Dollars (< 2.50 USD per student). In addition, the choice of laser cutting as a manufacturing process permits the sharing of the required digital files to produce LeDNA, enabling FabLabs, maker spaces, hackerspaces, and schools to locally fabricate the toolkits and replace lost or damaged pieces. Local production facilitates dissemination in a way that would be unfeasible for other methodologies. Moreover, each kit can be used by many students for years after the initial investment, while wet lab approaches require the continuous acquisition of new reagents. Similar to noncomputational educational games developed for other biology topics, LeDNA allows students to collaborate and collectively build knowledge. It is therefore situated in an intermediate zone between wet lab and virtual methodologies, combining the advantages of both approaches. While allowing hands-on, collective knowledge acquisition like laboratory activities, it does not require expensive and complex infrastructure. Furthermore, our results point out that it could be successfully combined with one of the many wet-lab approaches available to further increase its impact in resource-rich regions, while still producing significant results when applied alone in areas where those resources are not available.

While initially designed for teaching CRISPR-Cas, LeDNA can be expanded in the future to cover various topics in Molecular Biology. For example, the RNA ribonucleotides could be used to teach RNA interference, while the amino acids could be employed to teach the structure and mechanisms of action of proteins and enzymes. The open-access and modular nature of LeDNA enables researchers and educators worldwide to develop, implement, and share these and other modules that we cannot yet foresee.

## Supporting information

Material and methods, supplementary information and results, learning assessment questionnaires, and the design of each LeDNA part

Scalable Vector Graphics file for manufacturing each LeDNA toolkit piece

## SUPPORTING INFORMATION

Supplementary Word file containing material and methods, supplementary information, supplementary results, learning assessment questionnaires, and a description of the design of each LeDNA toolkit piece.

Supplementary Scalable Vector Graphics file for manufacturing each LeDNA toolkit piece (available in English, French, German, Portuguese, Spanish, and Urdu).

### Ethics approval

The study was approved by the Research Ethics Committee (CAAE: 55768021.0.0000.5426).

### Consent to participate

Participants and their legal guardians received Informed Consent and Assent Forms.

## Author Contribution

Kundlatsch conceived and designed the study, produced the material, and tested the protocols. Rodrigues designed the toolkit’s wooden case and provided technical assistance for using the laser cutting machine. Zocca supervised the plasmid construction and the wet lab protocol development. Kundlatsch, Zocca, Amorim, Paiva, and Neto graded the tests and compiled the data. Campos and Kundlatsch performed the statistical analysis. Pedrolli supervised the work and acquired funding. Kundlatsch and Pedrolli analyzed the data and wrote the first manuscript draft. All authors read and approved the final manuscript.

## ACKNOWLEDGEMENTS

We are thankful to the high school students of the *Escola Estadual - Professor Sebastião de Oliveira Rocha*, the science teacher Isabel Cristina Santana Kakuda and the director Lucinei Aparecida Tavoni Bueno. We also thank the undergrad and graduate students from Unesp. We extend our gratitude to Hassnain Qasim Bokhari for assisting in the translation of LeDNA into Urdu. Pedrolli’s research group is supported by the National Council for Scientific and Technological Development (CNPq) [grants 405490/2021-6 and 305324/2023-3].

