## Supplementary material for "LeDNA: a cut-and-build toolkit to democratize education on CRISPR gene editing technology": Material and methods, supplementary information and results, learning assessment questionnaires, and the design of each LeDNA part

**MATERIALS & METHODS**

**Design and Manufacturing of the Toolkit**

LeDNA was designed using Inkscape (1.0.2) and manufactured using Medium-density fiberboard (MDF) rectangular plates (300 mm x 200 mm x 3 mm), with 18 plates per kit and 7 per storage box. Pieces were cut on a 40W laser cutting machine (SL-320, CNC Zone), with trajectory instructions sent via CorelLaser. Cutting was performed at 40% power with a speed of 10 mm/s, while engraving utilized 10% power at a speed of 20 mm/s (using the cut function, resulting in a “half-cut”).

**Strains and plasmid construction**

The pEduc plasmid was constructed based on the pJOE8999^10^ by inserting the guide RNA targeting the GFP gene using the *Bsa*I restriction sites, and the donor DNA using the *Sfi*I restriction sites. The donor DNA is formed by the RFP gene and the homologous arms assembled from the PCR products through Golden Gate using *Bsa*I restriction sites (Figure S12 and Appendix 3). *E. coli* Top10 was used for cloning and plasmid propagation. The constructed plasmid was then used to transform *E. coli* JM109, which was used as plasmid source for *B. subtilis* transformation.

**Wet Lab Lecture**

We developed a simplified educational wet lab protocol based on the single-plasmid system for *B. subtilis*^10^. Some pre-work was required to be carried out in the university lab, such as competent cells preparation and plasmid construction (donor DNA and gRNA cloning). To enhance its applicability across diverse educational scenarios, the protocol was designed to be completed within a timeframe of 90 minutes or less. Moreover, it requires minimal on-site equipment, only a portable mini transilluminator (Figure S20). The students received frozen *B. subtilis* competent cells from a strain carrying a GFP gene integrated into the *amyE* locus in the genome^2^, along with the pJOE8999 plasmid (Figure S21) containing the RFP sequence as donor DNA and the machinery required to replace the former with the latter. The transformation protocol followed the Two-step *Bacillus subtilis* Transformation Procedure^3^. Starting the procedure, the students thawed the competent cells by holding the microtube tightly with their hands for one minute. Then, they transferred the pre-aliquoted SpII + EGTA medium to the microtube containing the competent cells using a disposable Pasteur pipette and mixed gently. The plasmid solution was next added to the tube in the same manner. After transformation, cells were plated on LB medium enriched with 5 µg/mL chloramphenicol and 0.5% (m/v) mannitol, and incubated on the bench at room temperature for 30 minutes. In the meantime, the participants received cells that had already undergone transformation and transferred them into an antibiotic-free medium to simulate the plasmid loss process. In the next step, they received another LB plate containing cells grown without plasmid and inoculated them into a fresh LB plate. Next, the students plated the cells they had transformed in the first step. All steps were carried out on the regular lab bench without any sterilization method other than applying 70% (v/v) ethanol to the surfaces. Finally, the students visualized the fluorescence of cells producing GFP and RFP using a portable mini transilluminator. The results of their transformations were presented to them the following week.

**Theoretical Lecture**

We developed a theoretical class consisting of three 30-minute blocks. The first block focused on presenting applications of the CRISPR technology relevant to the participants' daily lives, including its use in livestock^4,5^, agriculture^6,7^, combating disease-transmitting insects^8^, and human gene editing^9^. The second block reviewed topics previously covered in secondary school, such as DNA, RNA, and protein structure and properties, as well as transcription and translation mechanisms. Lastly, the final block aimed to explain the mechanism of the CRISPR-Cas technology, including the cellular components and processes required for its application by researchers.

**3-Stages Teaching Approach**

A 3-Stages Teaching Approach was implemented encompassing the LeDNA, wet lab, and theoretical lectures. Each intervention lasted 90 minutes and was applied at a week interval from each other (Figure 1). The three interventions were evaluated both individually and combined in different orders.

**Development of a methodology for learning assessment**

To assess the students' interest in biology, a questionnaire composed of five questions was adapted from Kennedy et al^11^ (Appendix 1). To measure students' learning outcomes a new tool was developed composed of three distinct sessions, each consisting of three questions. The first session assesses knowledge of basic genetics that are already included in the high school curriculum in Brazil. The second session assesses the technical aspects of how the CRISPR-Cas technology works. Finally, the third session assesses students' understanding of the technology's present and future impacts. Four different versions of the questions with minor changes were developed to reduce the interference in the test results of both the successive repetitions and the possibility of sharing answers between the participants. The different questions composing the learning assessment tool were developed to allow students to express knowledge in different ways. For this reason, the scoring system differs between them (Appendix 2). To minimize the effect of guessing, the students were informed that the test results would not affect their overall school scores. Additionally, in every question, the participants had the option of selecting "I don't know/I’d like not to answer." Both the interest and evaluation questionnaires were applied before any intervention and after each of them. The students also answered a socioeconomic questionnaire before the first intervention.

**Statistical analysis**

The test scores were adjusted to a Two-Parameter Logistic Model (2PL) of the Item Response Theory using jMetrik^12^ software to evaluate the developed assessment instrument. The parameters of difficulty and discrepancy were calculated for each of the questions that fit the model. Finally, the alpha coefficient (or Cronbach's Alpha)^13^ was calculated to assess the test reliability. The results from both the science attitude survey and the learning assessment tool were analyzed using JASP. First, the normality of the data distribution was assessed using the Shapiro-Wilk test. Next, Mauchly’s test of sphericity was performed. Finally, a repeated measures ANOVA was conducted, applying the Huynh-Feldt correction when sphericity was violated, followed by the Tukey test where applicable. The demographic data was evaluated using Chi-Square test or Fisher’s Exact test.

**Participants**

The evaluation of the different activities with high schoolers was conducted in collaboration with the *Escola Estadual de Ensino Integral Sebastião de Oliveira Rocha*, located in São Carlos, São Paulo, Brazil. All students enrolled in the institution's third year of high school in 2022 participated in the evaluation. The decision to select third-year students was agreed upon in consultation with the school's technical staff, considering the relevance of the topic to their class schedule and regular curriculum. A total of 90 students were divided into three groups of 30 and 56 of them participated in all sessions. The same evaluation was carried out with 68 undergraduate and graduate students from the São Paulo State University (UNESP), campus in Araraquara.

**Table S1.** CRISPR-Cas educational projects and the corresponding biological models applied

| **Biological System** | **Target Audience** | **Reference** |
| --- | --- | --- |
| *Arabidopsis thaliana* | Undergraduate students | Ruppel et al. ^14^ |
| *Caenorhabditis elegans* | Undergraduate students | Hastie et al. ^15^ |
| Cell-free | Undergraduate students | Collias et al. ^16^ |
| Cell-free | High School students | Stark et al. ^17^ |
| *Danio rerio* | Undergraduate students | Bhatt & Challa ^18^ |
| *Danio rerio* | Undergraduate students | Wolyniak et al. ^19^ |
| *Drosophila melanogaster* | Undergraduate students | Adame et al. ^20^ |
| *Escherichia coli* | Undergraduate students | Militello & Lazatin ^21^ |
| *Escherichia coli* | Undergraduate students | Pieczynski et al. ^22^ |
| *Escherichia coli* | High School students & non-scientists | Ziegler & Nellen ^23^ |
| Mammalian cell culture | Undergraduate students | Anderson ^24^ |
| *Saccharomyces cerevisiae* | Undergraduate & High School students | Sankaran et al. ^25^ |
| *Saccharomyces cerevisiae* | Undergraduate students | Sehgal et al^26^ |
| *Saccharomyces cerevisiae* | Undergraduate students | Vyas et al. ^27^ |
| *Saccharomyces cerevisiae* | Undergraduate students | Waal et al. ^28^ |
| *Saccharomyces cerevisiae* | Undergraduate students | Juríková et al. ^29^ |
| *Streptococcus thermophilus* | Undergraduate students | Hynes et al. ^30^ |
| *Vanessa cardui* | Undergraduate students | Thulluru et al. ^31^ |
| *Vanessa cardui* & *Xenopus laevis* | Undergraduate students | Martin et al. ^32^ |

**Table S2.** Projects applying board games to teach different biology topics

| **Subject** | **Reference** |
| --- | --- |
| Biochemistry | Rose et al. ^33^ |
| Biochemistry | Pennington et al. ^34^ |
| Cell Biology | Spiegel et al. ^35^ |
| Cell Biology | Carvalho et al. ^36^ |
| Evolution | Muell et al. ^37^ |
| Evolution | Miralles et al. ^38^ |
| Evolution | Luttikhuizen ^39^ |
| Genetics | Osier et al. ^40^ |
| Genetics | Osier et al. ^41^ |
| Immunology | Eckert et al. ^42^ |
| Immunology | Steinman et al. ^43^ |
| Microbiology | Coil et al. ^44^ |
| Molecular Biology | Barnes ^45^ |
| Nutrition | Amaro et al. ^46^ |
| Physiology | Chaves et al. ^47^ |

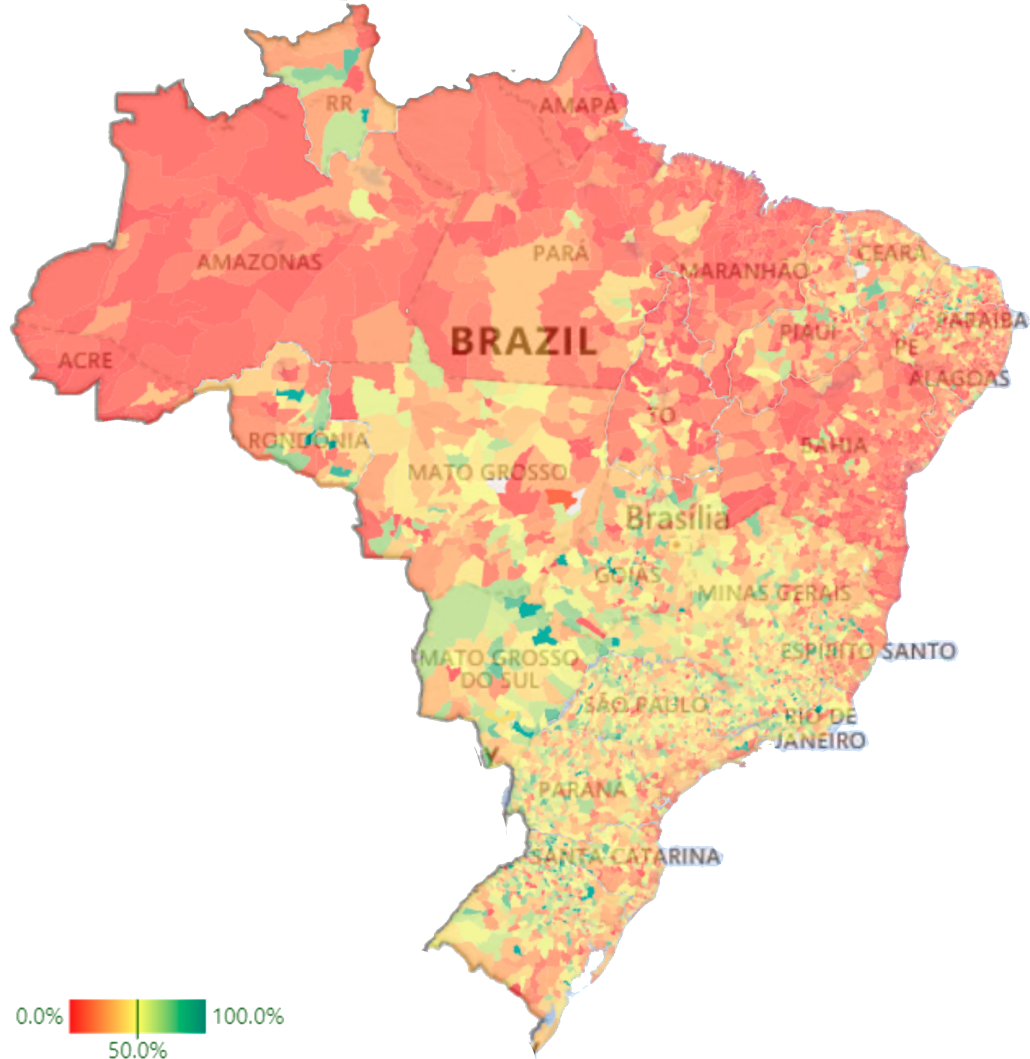

**Figure S1.** Distribution of schools with computer laboratories across municipalities in Brazil. The color of each municipality indicates the percentage of schools equipped with computers for student use. According to the 2023 School Census by the National Institute for Educational Studies and Research Anísio Teixeira (Inep), 30.2% of the 178,476 schools nationwide have dedicated computer laboratories^48^.

**Toolkit Design**

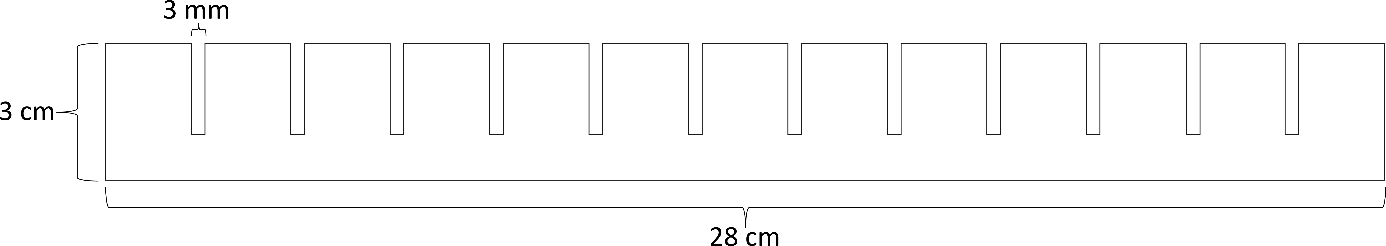

**Figure S2.** Sugar-phosphate backbone. A single piece was developed to represent the sugar-phosphate backbone of both double-stranded DNA and single-stranded RNA. It is 28 cm wide by 3 cm high, with grooves of 3 mm every 2 cm, allowing the attachment of 12 nucleotides. This number was chosen to allow the user to assemble a 3-amino acid polypeptide (the fourth codon should be a stop codon). The toolbox contains four identical copies of the part, two intended for the assembly of the DNA strand, one for the RNA strand, and one for the donor DNA for repair after the action of the Cas protein. For the construction of the guide RNA, a reduced version of the piece was developed, containing the same height and distance between the grooves, but allowing the attachment of only 6 nucleotides. The addition of nucleotides occurs perpendicularly.

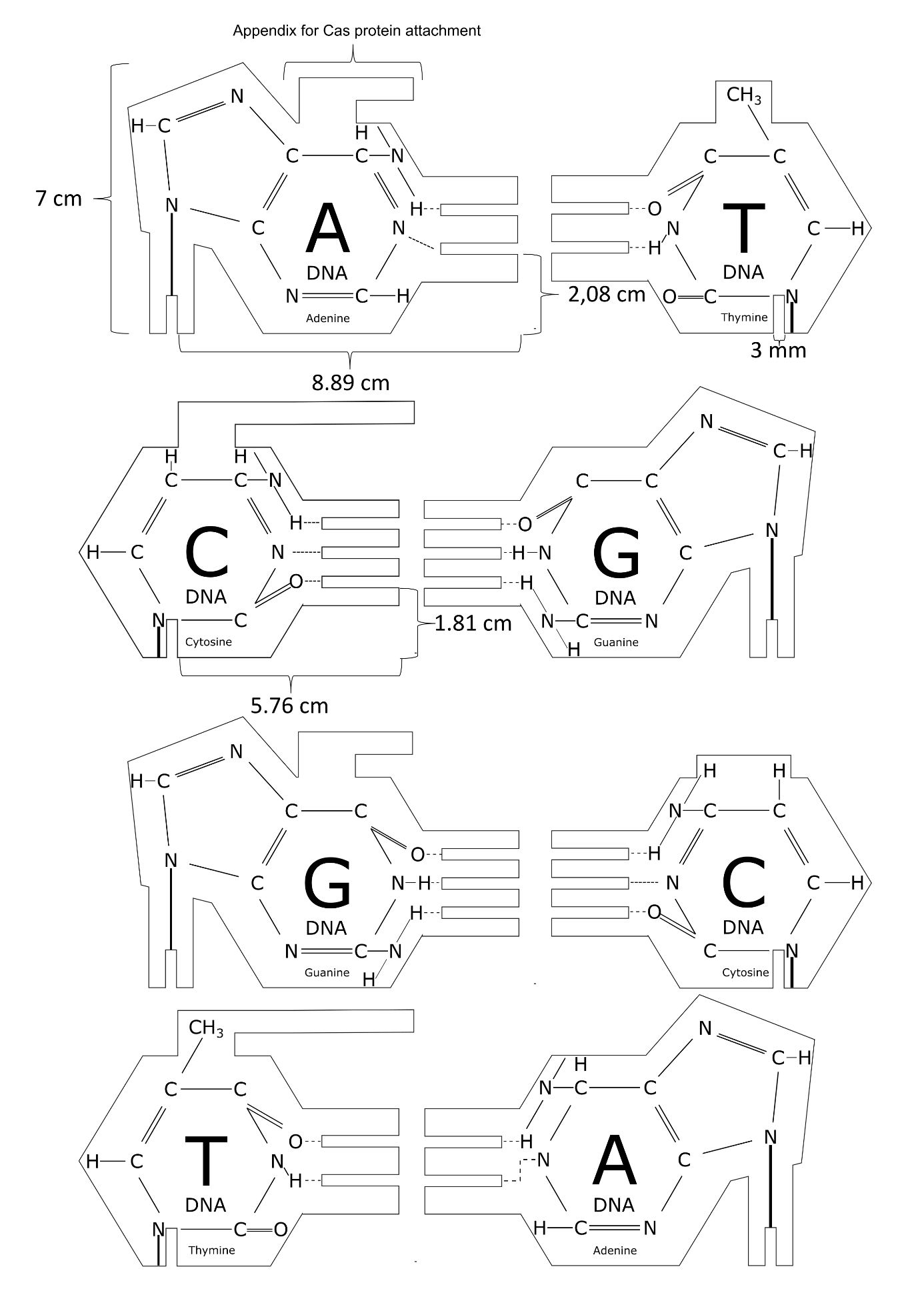

**Figure S3.** Double-stranded DNA. The connection of those pieces is made perpendicularly to the sugar backbone, with the hydrogen bonds shown in Figure S5 connecting the two DNA strands. The pieces used to assemble the template strand are different from those used to assemble the antisense strand. The complete kit contains 7 adenines, 7 cytosines, 5 guanines, and 8 thymines for the template strand, and 8 adenines, 5 cytosines, 7 guanines, and 7 thymines for the antisense strand. The number of each nucleotide is the maximum a student could use, considering that the first codon must be the start codon (ATG) and the last codon should be one of the three stop codons (TAA, TAG, and TGA). The molecular structure of the compound was added to the design of the piece, allowing the user to visualize the nucleotide. To facilitate its use, the name and an indication that it is a piece of DNA were also added.

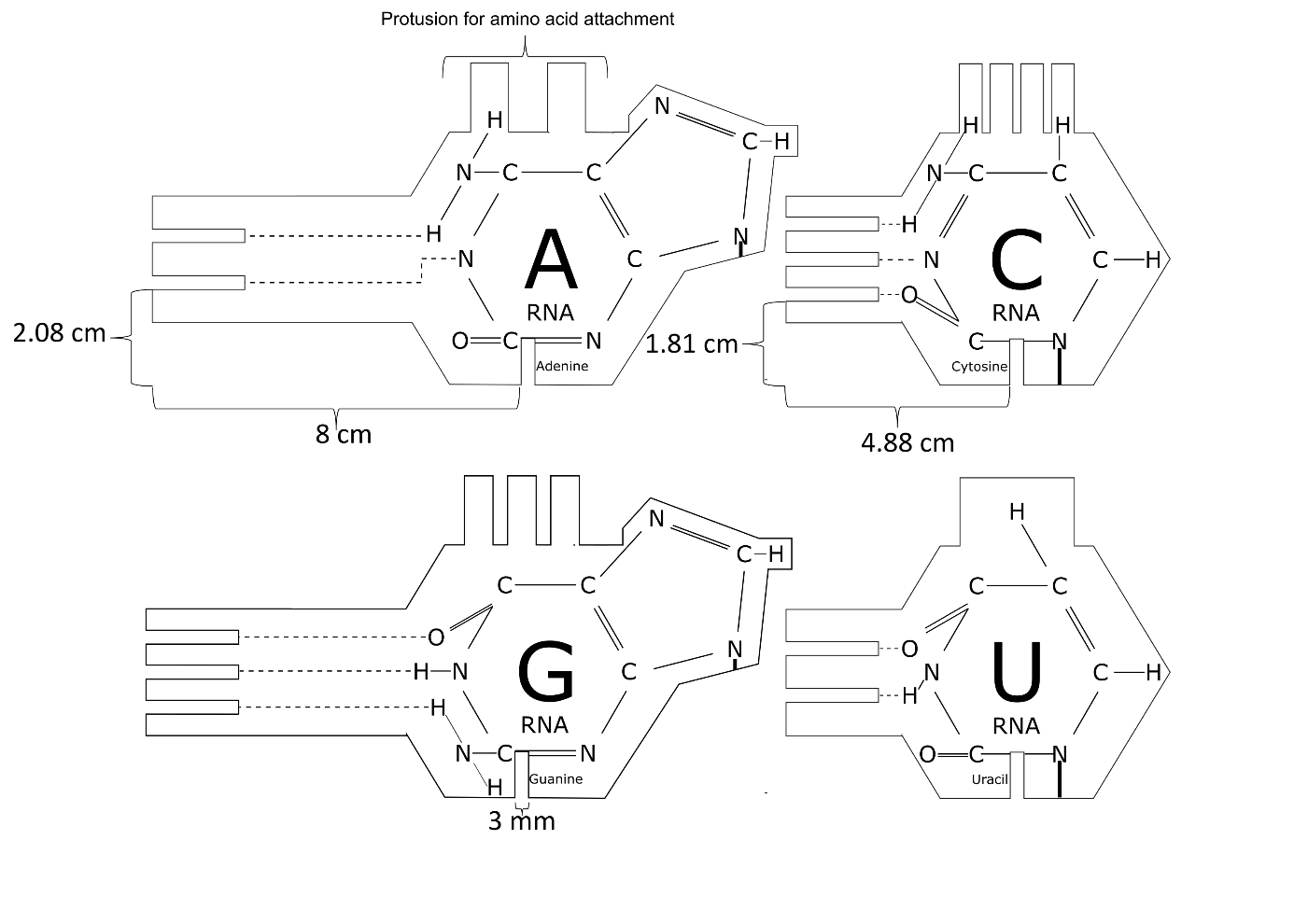

**Figure S4.** Single-stranded RNA. The connection is the same as the DNA nucleotides, perpendicularly to the sugar backbone. The complete kit contains 8 adenines, 5 cytosines, 7 guanines, and 7 uracils. Similarly to DNA nucleotides, part numbers differ due to start codon (AUG) and stop codon possibilities (UAA, UAG, and UGA). The chemical structure and name of each of the nucleotides were engraved. In addition, the term RNA was also included to facilitate the use of the toolkit.

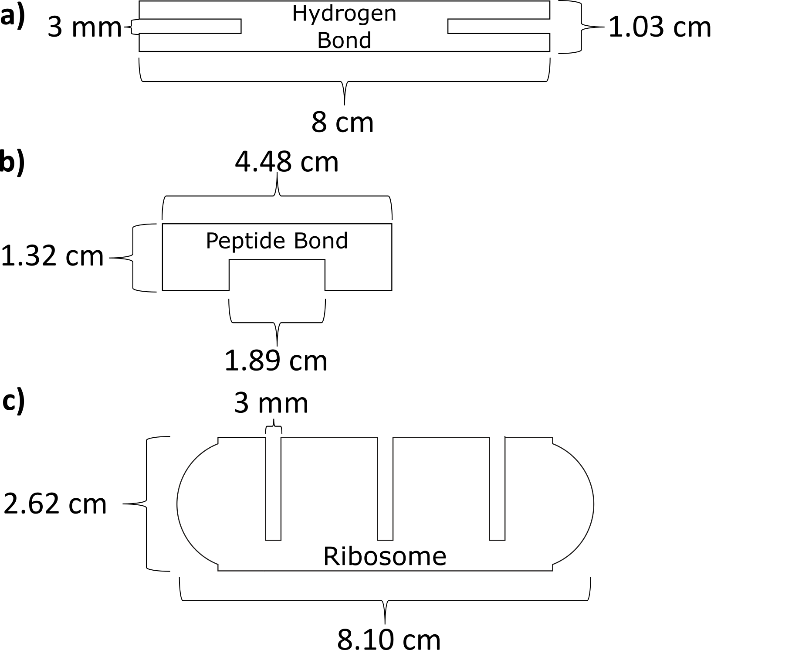

**Figure S5.** Hydrogen bond, peptide bond, and ribosome. a) Hydrogen bonds connect nucleotides. The kit contains 32 of them, the maximum number the user will need when assembling the DNA sequence: 7 for the start codon (ATG), 9 for the second and third codons (in case those are composed only of Cs and Gs), and 7 for the stop codon (in case TAG or TGA is used). b) Peptide bonds connect amino acids. Considering that the amino acid pieces are inserted over the already existing RNA strand, a different mechanism was employed. It is more accessible for the user to also insert the bond over the piece instead of laterally (as in the case of hydrogen bonds). The last of the four codons the user constructs must always be a stop codon. The piece representing the stop codon does not have a peptide bond slot precisely to stop the polypeptide formation. Frequently (except when the user adds a premature stop codon) three amino acids will be connected, therefore two bonds are needed to connect them. Four copies were added to the toolkit, allowing the formation of two polypeptides composed of three amino acids each. This allows the user to observe the polypeptide formed before and after the edition of the DNA sequence by CRISPR-Cas. c) The ribosome assembles the polypeptide. Its continuous connection gap allows it to fit any of the four nucleotides. The ribosome connects to three nucleotides (one codon) to facilitate accurate amino acid attachment and help teach the concept of codons to the user. Additionally, the side opening enables the piece to be pulled out from beneath the amino acid after connection, allowing it to be reused for the addition of the next amino acid.

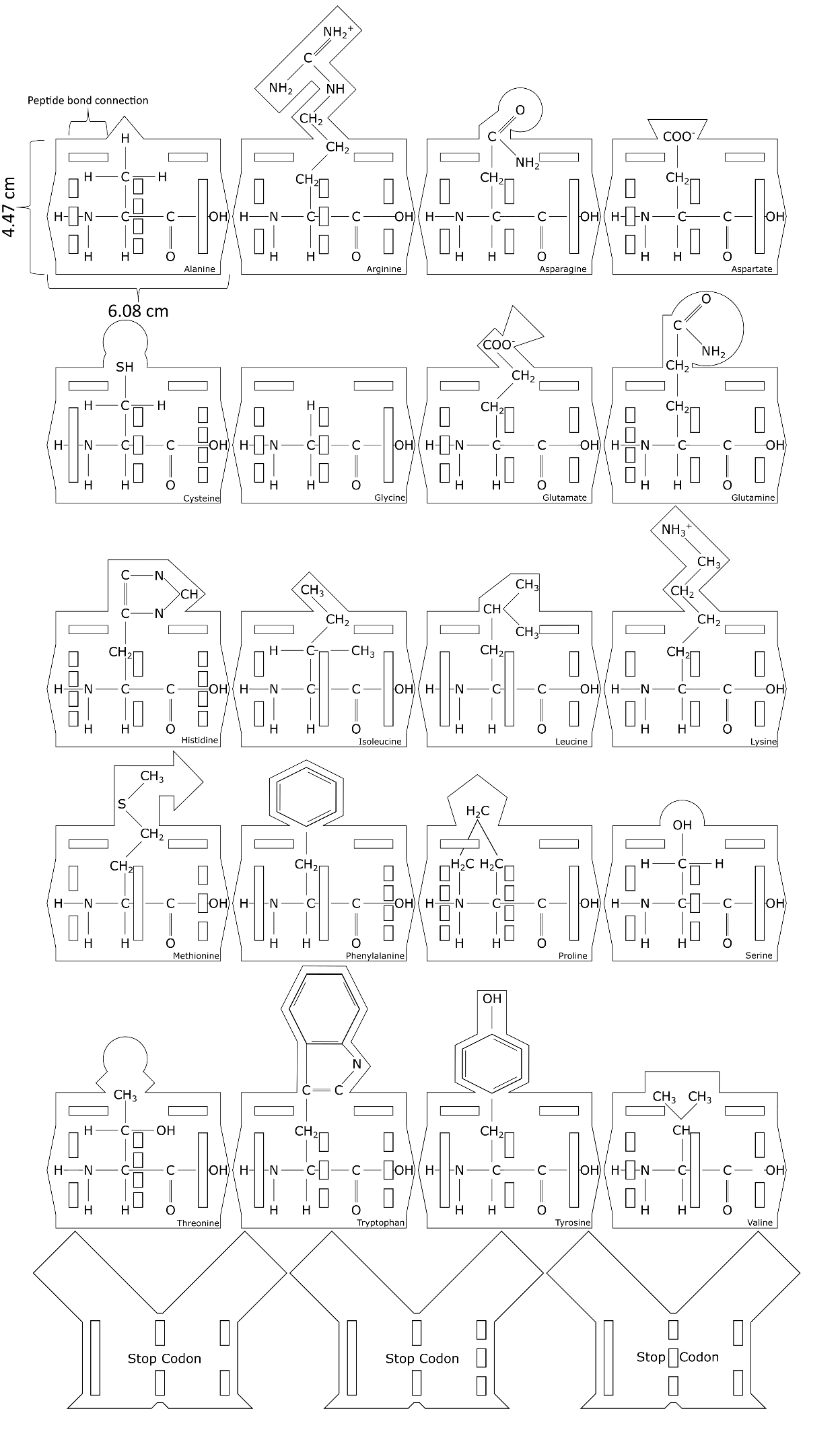

**Figure S6.** Amino acids and stop codons. For each of the twenty amino acids, a different design was developed, highlighting the different side chains and, therefore, their different physicochemical characteristics. Each amino acid also has a gap to connect with the peptide bond. In the case of the three stop codons, the gap for connecting with peptide bonds was removed, interrupting the translation process. A version of each amino acid was made for each codon that encodes it, keeping the exact side chain representation and adjusting to the proper cut pattern. The kit contains 64 pieces: alanine (4), arginine (6), asparagine (2), aspartate (2), cysteine (2), phenylalanine (2), glycine (4), glutamine (2), glutamate (2), histidine (2), isoleucine (3), lysine (2) leucine (6), methionine (1), proline (4), serine (6), tyrosine (2), threonine (4), tryptophan (1), valine (4), stop codon (3).

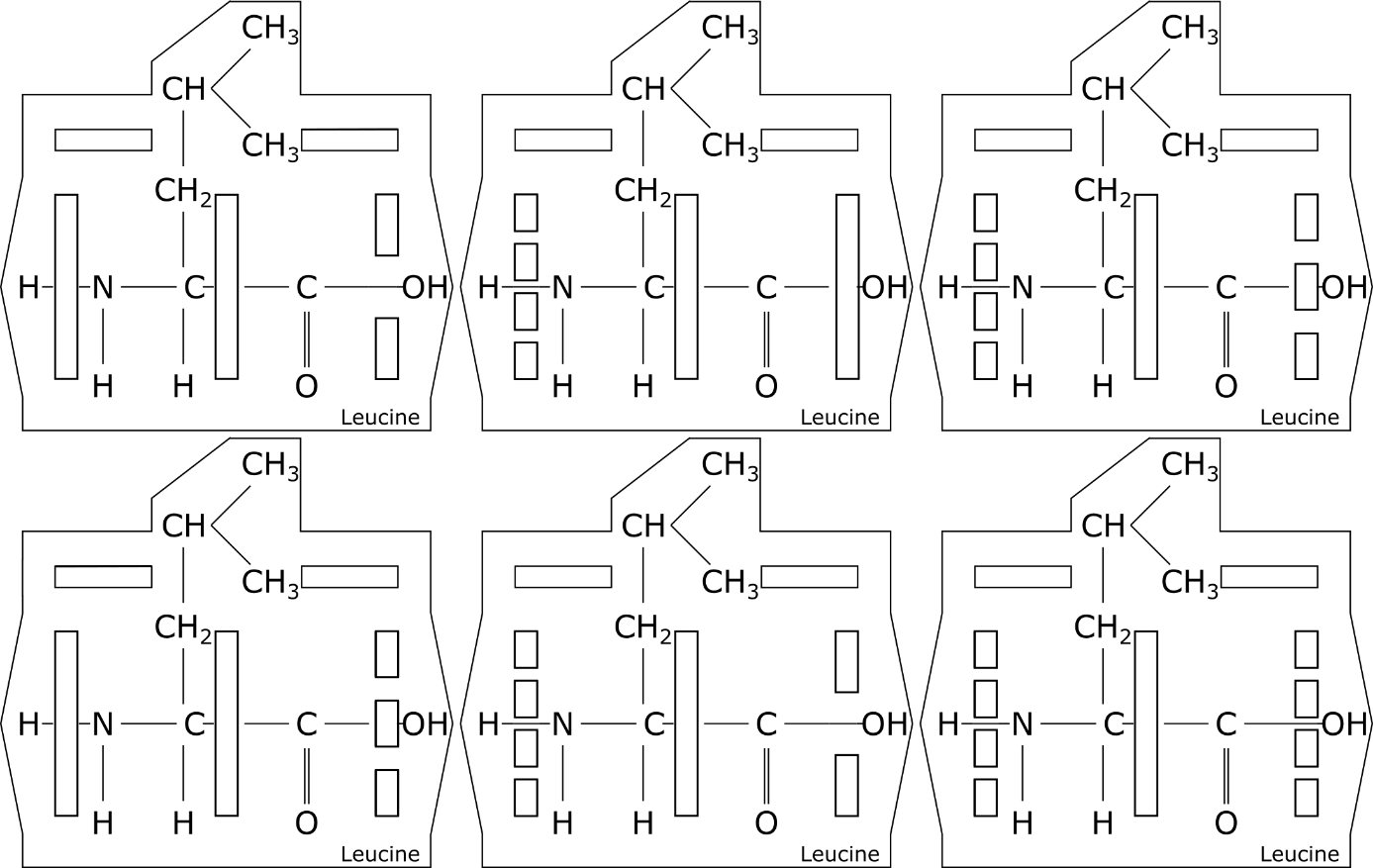

**Figure S7.** Different pieces representing leucine. Amino acids encoded by more than one codon were represented by similar pieces with modifications in the cut pattern. Only methionine and tryptophan are represented by a single piece in the kit.

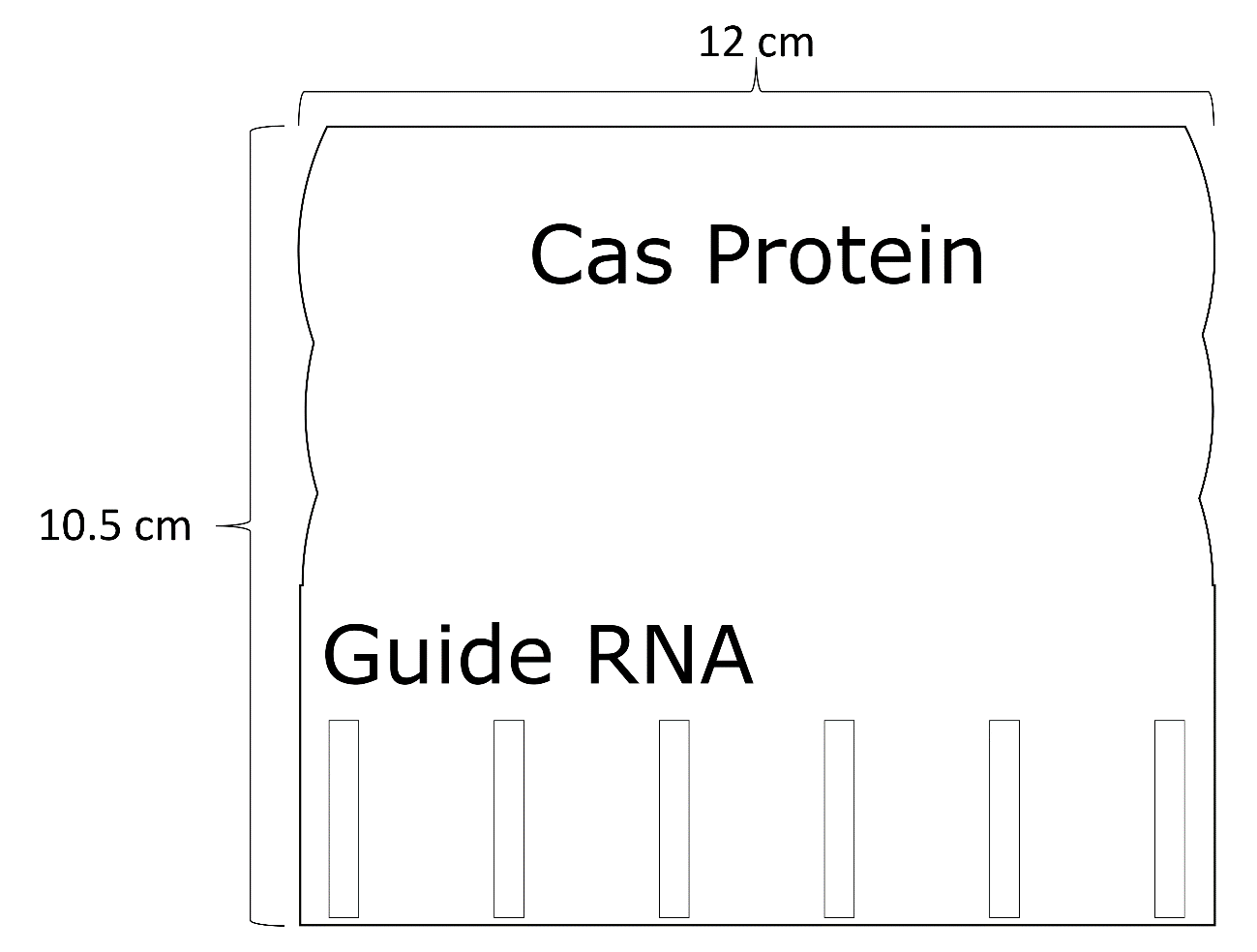

**Figure S8.** Cas protein and guide RNA. At the bottom edge of the part are the nucleotide slots that allow any RNA piece to be added. The top edge of the part is the point on the piece that connects under the appendix of the template DNA nucleotides and removes them when Cas is lifted by the user. The 6-nucleotide guide RNA is produced by the user by combining a shorter sugar-phosphate backbone with RNA nucleotides. The recognition and connection in the DNA strand occur through interaction with the guide RNA, followed by the cleavage of the DNA by the Cas protein. simulating the mechanism present in nature. After breaking the original double-stranded DNA, the user may use a DNA template for repair (Homology-Directed Repair) or insert nucleotides randomly (Non-Homologous End Joining).

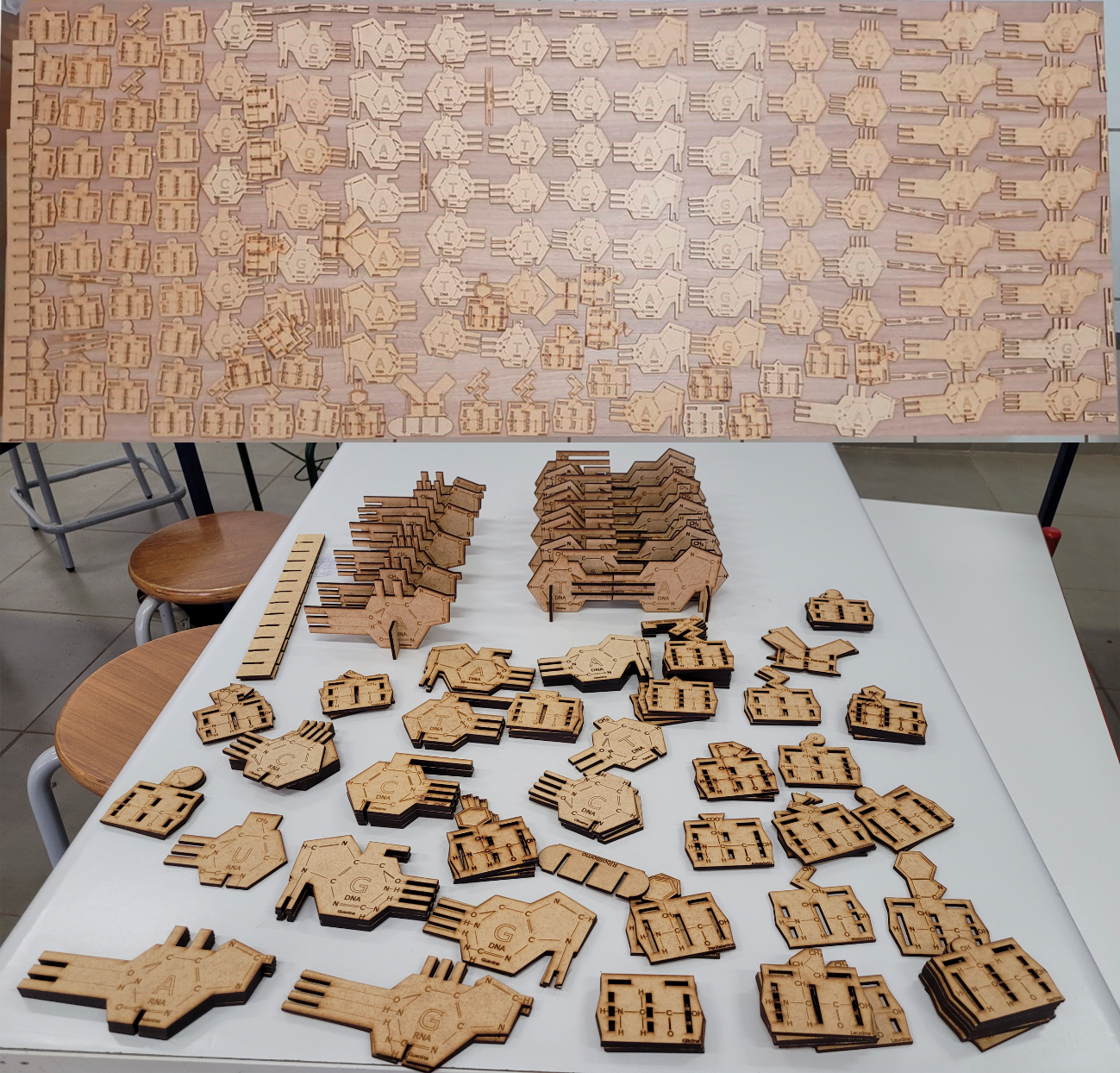

**Figure S9.** Complete LeDNA kit. At the top, each piece is presented individually. At the bottom, assembled DNA and RNA strands are presented and similar pieces are superimposed, to facilitate visualization. The complete set includes 27 nucleotides for template strand DNA, 27 for antisense strand DNA, 27 for RNA, 64 amino acids, 32 hydrogen bonds, 2 peptide bonds, 1 ribosome, 4 sugar-phosphate backbones, 1 sugar-phosphate backbone for guide RNA, 1 Cas protein.

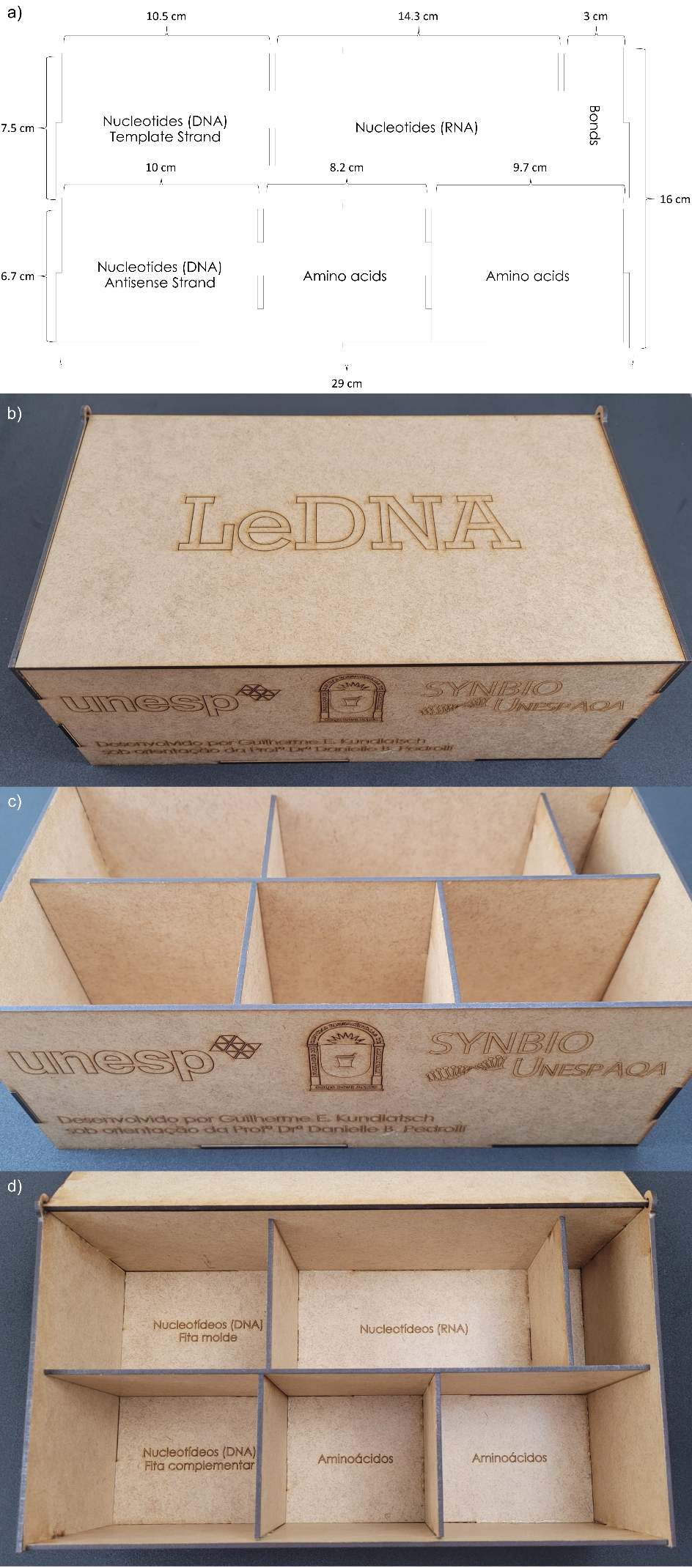

**Figure S10.** LeDNA storage box. The design of the base is represented in (a). The hollow regions between the compartments are used to fit the walls. The different compartments for amino acids have different measurements to accommodate the diversity of the size of the side chains of the pieces. On the lid (b) the name of the kit was added. On the front (c) the logos of the university and research group responsible for this project were engraved. The base (d) was built following the design shown in a).

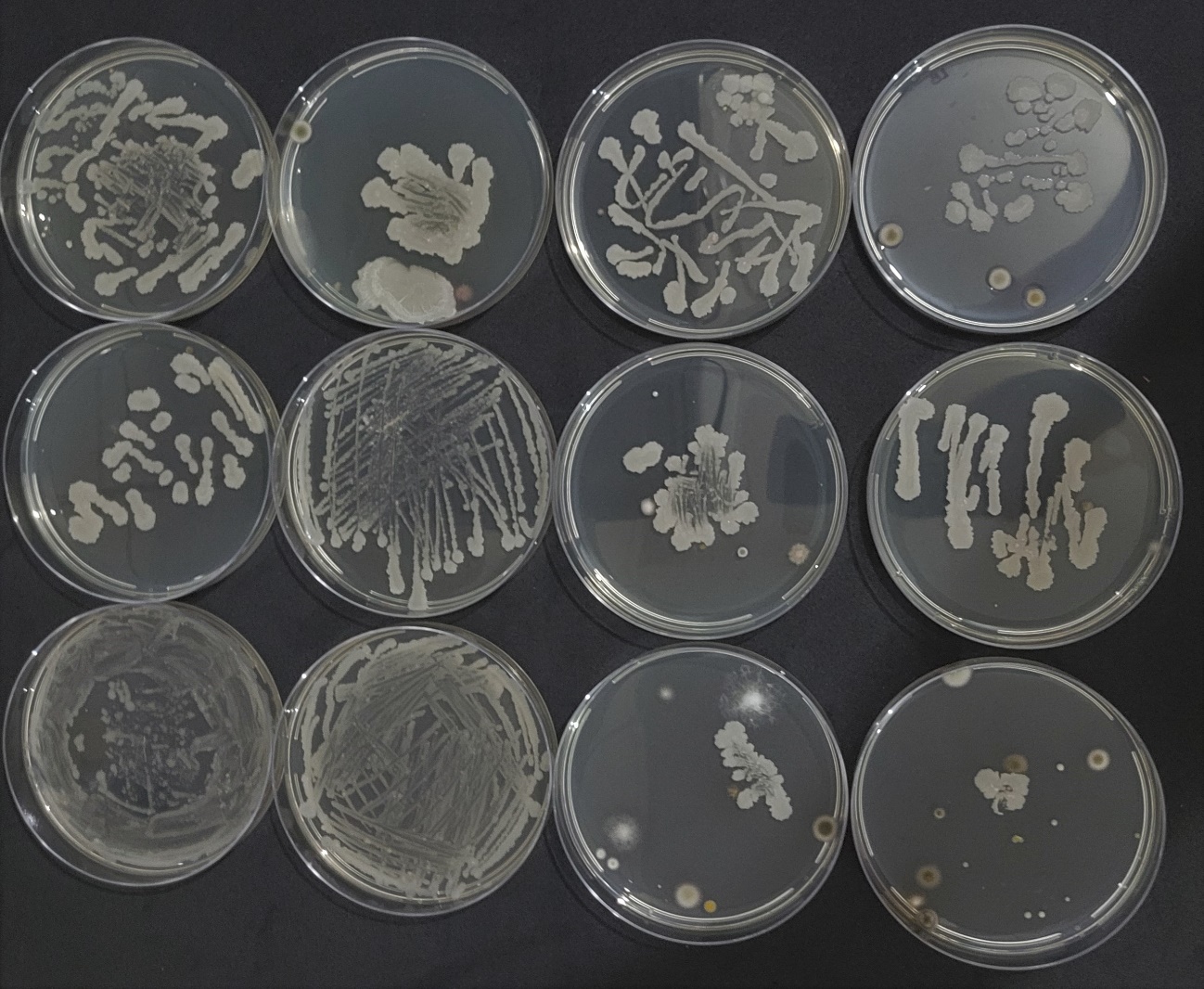

**Figure S11.** Results of *B. subtilis* transformations. Petri dishes streaked by students during the wet lab class showing white creamy colonies characteristic of *B. subtilis* across all plates. Additionally, five out of twelve plates exhibit yellow creamy colonies, as well as white and dark downy colonies, indicative of bacterial and fungal contamination, respectively.

**PARTICIPANTS’ DEMOGRAPHIC PROFILE**

Before the first intervention, a demographic assessment was conducted with each participant to identify possible differences between the three groups originally divided by their educational institution.

**Table S3.** Demographic profile of high school participants

|  | **Group 1** | **Group 2** | **Group 3** | **P*** |
| --- | --- | --- | --- | --- |
| Age | 17.00±0.535 | 17.07±0.583 | 17.04±0.367 | 0.897 |
| **Gender** | | | | |
| Male | 12 | 18 | 12 |  |
| Female | 9 | 11 | 14 |  |
| Other | 1 | 2 | 1 | 0.485 |
| **Monthly household income (in US dollars)** | | | | |
| < 400 USD | 7 | 6 | 9 |  |
| 400 – 800 USD | 8 | 9 | 8 |  |
| 800 – 2000 USD | 3 | 6 | 1 |  |
| 2000 – 4000 USD | - | 1 | 3 |  |
| Prefer not to answer | 4 | 9 | 7 | 0.325 |
| **Type of institution attended during elementary school (6 - 10 years)** | | | | |
| Public School | 20 | 23 | 24 |  |
| Private School | 2 | 3 | 3 |  |
| Both | - | 5 | 1 | 0.292 |
| **Type of institution attended during middle school (11 - 14 years)** | | | | |
| Public School | 21 | 24 | 25 |  |
| Private School | 1 | 1 | - |  |
| Both | - | 5 | 3 | 0.167 |
| **Educational level of participant's legal guardian** | | | | |
| No Education | - | - | 1 |  |
| Elementary school | 3 | 2 | 2 |  |
| Middle school | 3 | 6 | 5 |  |
| High school | 8 | 12 | 10 |  |
| Bachelor's degree | 4 | 5 | 3 |  |
| Postgraduate degree | 1 | 2 | 1 |  |
| Prefer not to answer | 3 | 4 | 6 | 0.992 |

**Note**. The categories “Other” and “Prefer not to answer” were not included in the analysis; *Chi-Square test or Fisher’s Exact test

**Table S4.** Demographic profile of undergraduate and graduate participants

|  | **Group 1** | **Group 2** | **Group 3** | **P*** |
| --- | --- | --- | --- | --- |
| Age | 19.97±1.468^a^ | 21.82±6.815^a^ | 26.93±4.42^b^ | <0.001 |
| **Gender** | | | | |
| Male | 10 | 10 | 7 |  |
| Female | 23 | 12 | 7 |  |
| Other | - | - | 1 | 0.342 |
| **Monthly household income (in US dollars)** | | | | |
| < 400 USD | 3 | 5 | 1 |  |
| 400 – 800 USD | 7 | 5 | 5 |  |
| 800 – 2000 USD | 6 | 9 | 6 |  |
| 2000 – 4000 USD | 10 | 2 | 1 |  |
| > 4000 USD | 1 | 1 | 1 |  |
| Prefer not to answer | 6 | - | 1 | 0.216 |
| **Type of institution attended during middle school (11 - 14 years)** | | | | |
| Public School | 9 | 12 | 7 |  |
| Private School | 22 | 7 | 5 |  |
| Both | 2 | 3 | 3 | 0.052 |
| **Type of institution attended during high school (15 - 17 years)** | | | | |
| Public School | 10 | 14 | 10 |  |
| Private School | 23 | 7 | 4 |  |
| Both | - | 1 | 1 | 0.005 |

**Note**. The categories “Other” and “Prefer not to answer” were not included in the analysis; *Chi-Square test or Fisher’s Exact test

**Attitude towards science**

Although improving the participants’ attitude toward science is not the primary objective of LeDNA, we decided to investigate whether the use of this tool would have any impact, particularly among high school students. We adapted the instrument developed by Kennedy et al.^11^, which is capable of assessing multiple aspects with a minimal number of items, making it more suitable for repeated interventions such as those proposed in this study. We did not observe any statistically significant impact from any of the different activities. One hypothesis to explain this result is that the activities were conducted with students in their final year of high school, a time when most have already decided on their intended career path.

**Table S5.** Attitude towards science before and after the different activities

| **High School Students** | | | | | |
| --- | --- | --- | --- | --- | --- |
| **Group** | **Score** | **Before** | **Stage 1** | **Stage 2** | **Stage 3** |
| **A** | Mean ± SD | 3.444±0.725 ^a, A^ | 3.544±0.745 ^a, A^ | 3.400±0.726 ^a, A^ | 3.544±0.757 ^a, A^ |
| **B** | Mean ± SD | 3.448±0.748 ^a, A^ | 3.438±0.755 ^a, A^ | 3.333±0.608 ^a, A^ | 3.457±0.738 ^a, A^ |
| **C** | Mean ± SD | 3.365±0.465 ^a, A^ | 3.471±0.474 ^a, A^ | 3.482±0.413 ^a, A^ | 3.541±0.504 ^a, A^ |
| **Undergraduate and Graduate Students** | | | | | |
| **Group** | **Score** | **Before** | **Stage 1** | **Stage 2** | **Stage 3** |
| **A** | Mean ± SD | 4.139±0.348 ^a, A^ | 4.127±0.367 ^a, A^ | 4.067±0.366 ^a, A^ | 4.103±0.407 ^a, A^ |
| **B** | Mean ± SD | 4.430±0.385 ^a, A^ | 4.590±0.370 ^a, A^ | 4.550±0.343 ^a, A^ | 4.400±0.395 ^a, A^ |
| **C** | Mean ± SD | 4.414±0.447 ^a, A^ | 4.357±0.465 ^a, A^ | 4.443±0.452 ^a, A^ | 4.429±0.407 ^a, A^ |
| LeDNA Wet Lab Theoretical | | | | | |

Uppercase letters indicate significant statistical differences between columns, while lowercase letters indicate significant statistical differences between rows.

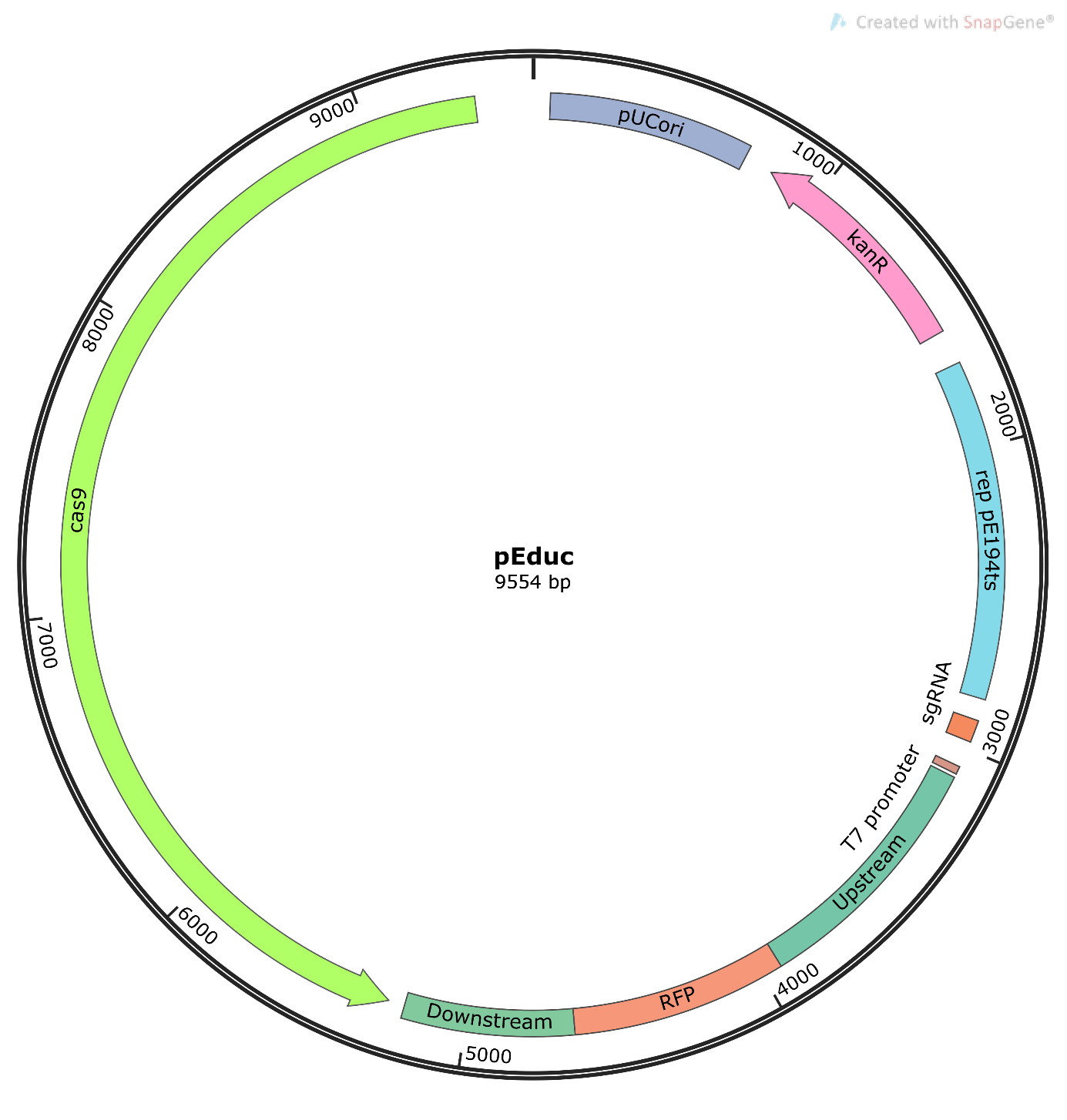

**Figure S12.** Map of the pEduc plasmid including the *cas9* coding sequence, sgRNA and donor DNA.

**Statistical validation of the test**

To assess students' interest in science, a series of previously validated instruments were available in the literature. However, a similar instrument was not available for the assessment of CRISPR-Cas learning. Consequently, we developed a novel instrument consisting of nine questions organized into three sections, each containing three questions. These sections were designed to assess participants' knowledge of Basic Genetics, CRISPR-Cas Mechanism, and CRISPR-Cas Applications. Responses from high school students who participated in all activities were accounted for, totaling 224 responses. The test scores were adjusted to a Two-Parameter Logistic Model (2PL) of the Item Response Theory (Table S3). The questions in the first section “Basic Genetics” resulted in a higher frequency of correct answers compared to the other two (Table S4) and presented a higher difficulty parameter (Table S5), which indicates that they are easier questions. This outcome was expected, as they assessed topics that students had previously studied in High School.

**Table S6.** Item fit to Two-Parameter Logistic Item Response Model. A p-value larger than 0.05 indicates that the question fits the model.

| **Question** | **p-value** |
| --- | --- |
| 1 | 0.4168 |
| 2 | 0.1702 |
| 3 | 0.8370 |
| 4 | 0.4890 |
| 5 | 0.5943 |
| 6 | 0.4426 |
| 7 | 0.2616 |
| 8 | 0.2111 |
| 9 | 0.9142 |

**Table S7.** Analysis of the frequency of responses to the evaluation form. Questions 1, 2, and 3 compose the “Basic Genetics” section, questions 4, 5, and 6 the “CRISPR-Cas Mechanism” section, and questions 7, 8, and 9, the “CRISPR-Cas Applications” session.

| **Question** | **Score** | **Frequency** | **Percentage** |
| --- | --- | --- | --- |
| 1 | 0 | 93 | 41.52% |
|  | 0.5 | 27 | 12.05% |
|  | 1 | 104 | 46.43% |
| 2 | 0 | 84 | 37.5% |
|  | 0.2 | 17 | 7.59% |
|  | 0.4 | 4 | 1.79% |
|  | 0.6 | 26 | 11.61% |
|  | 0.8 | 0 | 0% |
|  | 1 | 93 | 41.52% |
| 3 | 0 | 37 | 16.52% |
|  | 0.5 | 5 | 2.23% |
|  | 1 | 182 | 81.25% |
| 4 | 0 | 99 | 44.20% |
|  | 0.33 | 34 | 15.18% |
|  | 0.66 | 38 | 16,96% |
|  | 1 | 53 | 23.66% |
| 5 | 0 | 150 | 66.96% |
|  | 0.5 | 24 | 10.71% |
|  | 1 | 50 | 22.32% |
| 6 | 0 | 192 | 85.71% |
|  | 0.5 | 8 | 3.58% |
|  | 1 | 24 | 10.71% |
| 7 | 0 | 100 | 44.64% |
|  | 0.33 | 39 | 17.41% |
|  | 0.66 | 62 | 27.68% |
|  | 1 | 23 | 10.28% |
| 8 | 0 | 166 | 74.11% |
|  | 1 | 58 | 25.89% |
| 9 | 0 | 171 | 76.34% |
|  | 1 | 53 | 23.66% |

**Table S8.** Calculated difficulty and discrimination of each test item. The closest the Difficulty parameter is to 1, the easiest the question.

| **Question** | **Difficulty** | **Discrimination** |
| --- | --- | --- |
| 1 | 0.5848 | 0.5688 |
| 2 | 0.5071 | 0.6486 |
| 3 | 0.8237 | 0.4719 |
| 4 | 0.3957 | 0.6030 |
| 5 | 0.2768 | 0.5246 |
| 6 | 0.1250 | 0.4792 |
| 7 | 0.3428 | 0.7486 |
| 8 | 0.2589 | 0.5279 |
| 9 | 0.2366 | 0.5326 |

**Table S9.** High School participants' perception of the impact of CRISPR-Cas on their future careers. To illustrate the participants' perception of CRISPR-Cas at the end of the activities, some of the participants' responses to question 9 are listed: “Imagine that you travel 35 years into the future and find yourself. What do you think your profession will be? Will it be affected by CRISPR-Cas technology?”

| **Group** | **Answer** | **Career** |
| --- | --- | --- |
| Group A | “Modifying DNA to make muscle development easier through physical training” | Physical education |
| Group A | “Making students think faster to increase learning” | Pedagogy |
| Group A | “Everyone's daily life will be modified by CRISPR-Cas, because it can change the way in which any product is produced” | Computer engineering |
| Group A | “I will promote the sale of products developed with CRISPR technology” | Advertising and marketing |
| Group B | “Class materials (books) could be produced differently” | Social sciences |
| Group B | “In the movie Winter the Dolphin, the character loses his tail and has to wear a prosthesis so as not to harm his spine, in the future with CRISPR-Cas I could modify his DNA so that the tail could grow again” | Marine biology |
| Group C | “Clothing made by microorganisms modified with CRISPR-Cas can affect my work” | Dance |

**APPENDIX 1**

**Science attitude questionnaire**

This instrument was adapted from Kennedy et al^11^. Four different versions were developed with small modifications and randomly distributed.

Mark an X in the option (number or word) that best suits you.

1 – It is very likely that I will continue studying biology after finishing high school.

| Completely Agree |  | Partially Agree |  | Neither Agree Nor Disagree |  | Partially Disagree |  | Completely Disagree |
| --- | --- | --- | --- | --- | --- | --- | --- | --- |

2 – I think studying biology is

| Boring |  |  |  |  |  |  |  | Interesting |
| --- | --- | --- | --- | --- | --- | --- | --- | --- |
| 1 |  | 2 |  | 3 |  | 4 |  | 5 |

3 – I have difficulty completing biology activities.

| Completely Agree |  | Partially Agree |  | Neither Agree Nor Disagree |  | Partially Disagree |  | Completely Disagree |
| --- | --- | --- | --- | --- | --- | --- | --- | --- |

4 – I think I'm pretty good at biology.

| Completely Agree |  | Partially Agree |  | Neither Agree Nor Disagree |  | Partially Disagree |  | Completely Disagree |
| --- | --- | --- | --- | --- | --- | --- | --- | --- |

5 – Working as a scientist would be interesting.

| Completely Agree |  | Partially Agree |  | Neither Agree Nor Disagree |  | Partially Disagree |  | Completely Disagree |
| --- | --- | --- | --- | --- | --- | --- | --- | --- |

**APPENDIX 2**

**CRISPR-Cas Learning assessment questionnaire**

Four distinct versions of the questionnaire were developed, featuring varied sequences of options and DNA sequences, which were then randomly distributed. This aimed to mitigate both cheating tendencies and the likelihood of participants memorizing specific patterns. Furthermore, it was explicitly communicated to the participants that their responses would remain anonymous and would not bear any impact on their academic records. This approach was adopted to diminish the potential influence of guesswork on the accuracy of their learning assessments. Finally, the students received a codon table to assist them. They had 10 minutes to complete the questionnaire.

**Section 1. Basic Genetics**

**Question 1**

a) Which messenger RNA would be produced from this DNA sequence:

5’- TACGAGTAAATTGCA - 3’

3’- ATGCTCATTTAACGT - 5’

b) Which amino acids would be produced from the messenger RNA sequence generated in a)?

Scoring: The participant who writes the correct sequence of messenger RNA gets 0.5 points, getting another 0.5 if he describes the correct amino acids.

**Question 2**

Organize these terms into the appropriate boxes: Protein, RNA, DNA, Ribosome, RNA Polymerase.

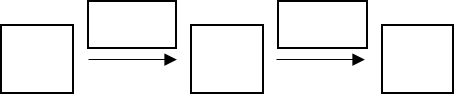

Scoring: For each correctly filled box, the participant receives 0.2 points.

**Question 3**

Draw a line connecting the nucleotides that pair in the formation of the double-stranded DNA.

Adenine	Adenine

Cytosine	Cytosine

Guanine	Guanine

Thymine	Thymine

Scoring: For each item correctly connected the participant receives 0.25 points

**Section 2. CRISP-Cas Mechanism**

**Question 4**

You are a scientist in your laboratory, and you would like to correct a defective gene in a particular organism. Mark with an X which items you need to insert into the organism for genome editing.

Ribosome	Mitochondria	Messenger RNA	guide RNA

CRISPR protein	Cas protein	Template DNA	mitochondrial DNA

Scoring: For each correct item the participant receives 0.33 points. For each incorrect item, the student loses 0.33 points, and the final score of the question cannot be less than 0.

**Question 5**

After the success of your experiment in the previous question, you would now like to disable another gene instead of modifying it. You repeat the previous steps, but this time you DO NOT include which of the previous items?

Ribosome	Mitochondria	Messenger RNA	guide RNA

CRISPR protein	Cas protein	Template DNA	mitochondrial DNA

Scoring: If the participant selects the correct answer, the student receives 1 point. If the participant selects the correct answer and any additional item, the student receives 0.5 points.

**Question 6**

You would like to replace the codon that encodes the amino acid ASPARTATE with the codon for GLUTAMATE in a given protein. You used CRISPR-Cas technology to cleave the genome, which DNA sequence would you use as a donor for the repair?

Protein-coding DNA :	Template strand 5’- ...GTACTGTCG... -3’

Antisense strand 5’- ...CATGACAGC... -3’

Scoring: The student who describes the correct donor DNA receives 1 point. The participant who describes a donor that also promotes changes other than the desired one receives 0.5 points.

**Section 3. CRISPR-Cas Applications**

**Question 7**

Mark activities for which CRISPR-Cas technology is already used/may be used in the future.

**Question 7**

Please mark activities for which CRISPR-Cas technology is already used/may be used in the future.

Modify embryonic cells to prevent hereditary diseases

Develop drought resistant plants

Create yeasts with greater biofuel production capacity

Chemical synthesis of new antibiotics

Scoring: For each correct item the participant receives 0.33 points. For each incorrect item the student loses 0.33 points.

**Question 8**

Briefly describe an activity in your daily life that could be impacted by CRISPR-Cas technology.

Scoring: In this open question, the participant receives 1 point for any answer, as long as it is justified.

**Question 9**

Imagine that you travel 30 years into the future and find yourself. What do you think your profession will be? Will it be affected by CRISPR-Cas technology? Justify briefly.

Scoring: In this open question, the participant receives 1 point for any answer, as long as it is justified.

**APPENDIX 3**

**DNA sequences used to construct pEduc**

**sgRNA:** GCTAGAAATAGCAAGTTAAAATAAGGCTAGTCCGTTATCAACTTGAAAAAGTGGCACCGAGTCGGTGCTTTTT

**Donor DNA:**

**Upstream homology region:** CTCCTGATCCAAACATGTAAGTACCAATAAGGTTATTTTTTAAATGTTTCCGAAGTATTTTTTTCACTTTATTAATTTGTTCGTATGTATTCAAATATATCCTCCTCACTATTTTGATTAGTACCTATTTTATATCCATAGTTGTTAATTAAATAAACTTAATTTAGTTTATTTATAGATTTCATTGGCTTCTAAATTTTTTATCTAGATAATAATTATTTTAGTTAATTTTATTCTAGATTATATATGATATGATCTTTCATTTCCATAAAACTAAAGTAAGTGTAAACCTATTCATTGTTTTAAAAATATCTCTTGCCAGTCACGTTACGTTATTAGTTATAGTTATTATAACATGTATTCACGAACGAAAATCGCCATTCGCCAGGGCTGCAGGAATTCGACTCTCTAGCTTGAGGCATCAAATAAAACGAAAGGCTCAGTCGAAAGACTGGGCCTTTCGTTTTATCTGTTGTTTGTCGGTGAACGCTCTCCTGAGTAGGACAAATCCGCCGCTCTAGCTAAGCAGAAGGCCATCCTGACGGATGGCCTTTTTGCGTTTCTACAAACTCTTGTTAACTCTAGAGCTGCCTGCCGCGTTTCGGTGATGAAGATCTTCCCGATGATTAATTAATTCAGAACGCTCGGTTGCCGCCGGGCGTTTTTTATGCAGCAATGGCAAGAACGTTGCTCGAGGGTAAATGTGAGCACTCACAATTCATTTTGCAAAAGTTGTTGACTTTATCTACAAGGTGTGGCATAATGTGTGTAATTGTGAGCGGATAACAATTAAGCTTAGTCGACAGCTAGCTGATTAACTAATAAGGAGGACAAAC

**Red Fluorescent Protein:** ATGGCTTCCTCCGAAGACGTTATCAAAGAGTTCATGCGTTTCAAAGTTCGTATGGAAGGTTCCGTTAACGGTCACGAGTTCGAAATCGAAGGTGAAGGTGAAGGTCGTCCGTACGAAGGTACCCAGACCGCTAAACTGAAAGTTACCAAAGGTGGTCCGCTGCCGTTCGCTTGGGACATCCTGTCCCCGCAGTTCCAGTACGGTTCCAAAGCTTACGTTAAACACCCGGCTGACATCCCGGACTACCTGAAACTGTCCTTCCCGGAAGGTTTCAAATGGGAACGTGTTATGAACTTCGAAGACGGTGGTGTTGTTACCGTTACCCAGGACTCCTCCCTGCAAGACGGTGAGTTCATCTACAAAGTTAAACTGCGTGGTACCAACTTCCCGTCCGACGGTCCGGTTATGCAGAAAAAAACCATGGGTTGGGAAGCTTCCACCGAACGTATGTACCCGGAAGACGGTGCTCTGAAAGGTGAAATCAAAATGCGTCTGAAACTGAAAGACGGTGGTCACTACGACGCTGAAGTTAAAACCACCTACATGGCTAAAAAACCGGTTCAGCTGCCGGGTGCTTACAAAACCGACATCAAACTGGACATCACCTCCCACAACGAAGACTACACCATCGTTGAACAGTACGAACGTGCTGAAGGTCGTCACTCCACCGGTGCTTAATAACGCTGATAGTGCTAGTGTAGATCGC

**Downstream homology region:**

CAGCTGCTGGTATTACACATGGTATGGATGAATTGTATAAATAATAATGAGCACTAGTCAAGGTCGGCAATTCTGCAGTACTAGGACGCCGCCAAGCCAGCTTAAACCCAGCTCAATGAGCTGGGTTTTTTGTTTGTTAAAAATGAAGAAGAAACTGTGAAGCGTATTTATAGCAAAGCACTCAAAAGTTTACCTTATGGGTGCTTTTTTCGTGCTTTTTTGAAAAGACAAAAAAAAGAACCTTGCCAAGCAAGATTCTTTGCATGCAAGCTAATTCGGTGGAAACGAGGTCATCATTTCCTTCCGAAAAAACGGTTGCATTTAAATCTTACATATGTAATACTTTCAAAGACTACATTTGTAAGATTTGATGTTTGAGTCGGCTGAAAGATCGTACGTACCAATTATTGTTTCGTGATTGTTCAAGCCATAACACTGTAGGGATAGTGGAAAGAGTGCTTCATCTGGTTACGATCAATCAAATATTCAAACGGAGGGAGACGATTTTGATGAAACCAGTAACGTTATACGATGTCGCAGAGTATGCCGGTGTCTCTTATCAGACCG
